# Rapid high-resolution size distribution analysis for adeno-associated virus using high speed SV-AUC

**DOI:** 10.1101/2023.05.08.539919

**Authors:** Steven A. Berkowitz, Nicholas Larson, George Bou-Assaf, Thomas Laue

## Abstract

When optimized, sedimentation velocity analytical ultracentrifugation (SV-AUC) provides the most-accurate, broadest-range, and highest-resolution size distribution analysis of any method. Generating simulated data for an adeno-associated virus (AAV) sample consisting of four species differing only in their DNA content and having closely spaced sedimentation coefficients, allows manipulation of the SV-AUC experimental protocol to optimize the size distribution resolution. In developing this high speed SV-AUC (hs-SV-AUC) protocol several experimental challenges must be overcome: 1) the need for rapid data acquisition, 2) avoiding optical artifacts from steep boundaries and 3) overcoming the increased potential for convection. A protocol, hs-SV-AUC, has been developed that uses high rotor speeds, interference detection and low temperatures to overcome these challenges. By confining data analysis to a limited radial-time window and using a very short run time (< 20 min after temperature equilibration), the need to match the sample and reference solvent composition and meniscus positions is relaxed, making interference detection is as simple to employ as absorbance detection. Experimental size distributions from the same AAV sample by hs-SV-AUC at 45K rpm and 10 °C versus low-speed SV-AUC at 15K rpm, and 10 °C illustrates the improved size distribution resolution offered by the hs-SV-AUC protocol.

## 1. Introduction

Sedimentation velocity analytical ultracentrifugation (SV-AUC) can provide the most-accurate, broadest-range, and highest-resolution particle size distribution analysis of any method [1–4]. The size distribution resolving power of SV-AUC is proportional to *rpm*^2^ · ℎ, where rpm is the rotor speed and *h* is the radial distance between the sample meniscus and the cell base [5–6]. For samples having components with sedimentation coefficients between ∼3-30S (at 20 °C), experimental protocols typically use rotor speeds between ∼30-50K rpm. In these cases, increasing the rotor speed to 60,000 rpm (the maximum rotor speed of the analytical ultracentrifuge) only improves resolution marginally, while reducing the number of samples per run. However, in the case of the large (> 30S), complex supramolecular biopharmaceuticals being developed for gene therapy (e.g. viral vectors and nanoparticles), rotor speeds typically do not exceed 20K rpm [7,8] and can be as low as 3K rpm [9]. Samples run at higher rotor speeds should provide a better-resolved sample size distribution. To date, little attention has been paid to optimizing experimental protocols for SV-AUC analysis of supramolecular biopharmaceuticals. Instead, from the authors’ perspective, the focus has been directed towards acquiring more data scans per sample in the hopes of improving the data analysis precision and maximizing the number of samples that can be run in an SV-AUC experiment. In order to acquire a large number of data scans, current SV-AUC protocols use lower rotor speeds and an increased run time. However, the longer run time allows diffusion to obscure boundary separation, decreasing the size distribution resolution of any analysis method, regardless of its numerical sophistication. Consequently, a balance must be struck between increased rotor speed, which improves boundary separation and reduces run time, and the time needed to acquire enough data scans per sample to allow effective data analysis.

Adeno-associated virus (AAV) is used as a model system to explore how increasing the rotor speed might improve SV-AUC size distribution analysis of gene therapy biopharmaceuticals. The choice of AAV is relevant given its widespread use in gene therapy [10–12], coupled with the need to characterize the wide range of drug-related impurities (e.g., empty, partially filled, over filled and aggregated AAV particles) commonly found in AAV preparations [13–16]. Due to diffusion, and the very heterogeneous nature of some of these impurities, size distribution analysis from normal low speed (10-20K rpm [7,8,13,17]) SV-AUC on AAV samples can result in overlapping and misleading size distributions. Consequently, AAV sample characterization may be compromised with respect to its homogeneity, the complexity of its heterogeneity, and the accurate quantification of the final AAV drug product potency. Uncertainty in these properties raises questions concerning the accuracy of the dosing solution being administrated to the patient.

An optimized high-speed SV-AUC (hs-SV-AUC) protocol for AAV initially was developed using simulations of a hypothetical AAV sample consisting of equal amounts of four homogeneous AAV capsids containing differing amounts of ssDNA in 0.15 M NaCl. The four AAV species differed in their sedimentation coefficients (at 20 °C) by a spacing of 3.7 Svedberg units, S, spanning a range of 89-100 S_20°_. Simulations at different rotor speeds show a significant resolution improvement is achieved by hs-SV-AUC over lower-speed protocols (10-20K rpm, which we have chosen to be represented by the average rotor speed of 15K rpm).

In addition to improved resolution, hs-SV-AUC results in very short (< 20 min) run times. These short run times coupled with the high molecular weight (MW) of AAV particles yields steep migrating boundaries due to their low diffusion. The presence of these steep, rapidly-moving boundaries mean that the optical detector must have high optical resolution, be insensitive to steep refractive gradients and provide rapid data acquisition. Absorbance detection is too slow (approximately in the range of 14-20 sec/cell for the Optima and 30-60 sec/cell for the XL-A/I depending on the radial increment and rotor speed), lacks radial resolution (50 μm) and is too sensitive to schlieren effects in steep boundaries for this application [18,19]. However, the radial resolution (9 μm), acquisition time (∼5 s) and accurate tracking of steep gradients of the interference (refractometric) detector (the Rayleigh interferometer, available and frequently also present on many analytical ultracentrifuges) meet the demands of hs-SV-AUC. Furthermore the interferometer captures the entire cell image simultaneously (i.e., it does not scan the cell), so there is no data distortion due to the time lag between the acquisition of the first and last radial data points. The interferometer has vastly superior precision, orders of magnitude broader linear concentration response and far lower sensitivity to light scattering and schlieren distortion compared to absorbance detection.

Nevertheless, interferometers are sensitive to the net differential movement of *all* solution components in both the reference and sample sectors. Ordinarily, to prevent solvent component sedimentation from contributing to the interference signal: 1) the solvent composition of components must be matched exactly in the sample and reference sectors and 2) the sample and reference menisci positions also must be matched, making cell filling onerous or necessitating the use of delicate meniscus matching centerpieces. Software has been added to SEDFIT in an attempt to mitigate situations where these conditions are not achieved [20]. It also has been proposed that water, rather than salt, be used as the reference material to avoid mismatch. However, there are two caveats to using just water. First, the quality of the resulting interference fringes depends on the light source coherence length, which is equal to the laser center wavelength squared divided by the laser bandwidth. The authors’ experience is that the coherence length of laser diodes can be quite short, varies with operating conditions, and varies significantly from diode to diode, resulting in washed-out fringes and poor data quality. Second, using water will not eliminate the salt contribution to the signal. Instead, the entire diffraction envelop will be shifted upwards, and the boundary from the salt sedimentation will dominate the signal, complicating any subsequent analysis.

Here we show that in both computer simulation and actual experiments that by restricting the run time and the radial range for data analysis (the “radial-time window”) the hs-SV-AUC protocol avoids contributions from the solvent boundary, resulting in a flat plateau region through which the AAV boundaries sediment. The excess plateau decrease due to solvent radial dilution is removed from the AAV signal during analysis with minimal loss in size distribution resolution.

However, the short run time precludes the establishment of a stabilizing solvent density gradient, so the hs-SV-AUC protocol is susceptible to convection distorting the boundaries, especially at high rotor speeds. To avoid the potential for convection, the hs-SV-AUC protocol should be conducted at a low temperature (e.g. ≤ 10 °C) which takes advantage of the unique temperature dependent density properties of water [21]. In addition, the lower temperature also results in slower boundary movement, allowing for a longer running time and the acquisition of more data.

Finally, AAV samples in formulation buffer were analyzed using both a ‘standard’ low-speed protocol and the new hs-SV-AUC protocol. Results from these experiments provide direct experimental evidence that the AAV hs-SV-AUC protocol results in a significantly higher resolution size distribution analysis. An assessment of the advantages, limitations, and options for overcoming these limitations are discussed.

## 2. Material & Methods

### 2.1 AAV and other material

All AAV-AUC work was carried out using the same AAV sample produced, purified and diafiltered into an AAV formulation buffer at Biogen. AAV stock solutions were provided at concentrations requiring a 10- to 20-fold dilution with buffer to generate two different AAV sample concentrations. For solvent mismatch SV-AUC experiments, either 150 mM NaCl, PBS (137 mM NaCl, 2.7 mM KCl, 4.3 mM Na_2_HPO_4_, pH 7.4) or AAV formulation buffer were used. The AAV formulation buffer used in this work has NaCl as the main component, along with low levels of a buffer and non-ionic detergent yielding a pH 7 solution.

#### 2.1.2 Hypothetical AAV sample

All simulations used a hypothetical AAV sample consisting of four species (Table 1): 1) the full AAV particle consists of a protein shell with an assigned MW of 3.90 MDa and a ssDNA genome (DNA payload) with an assigned MW of 1.33 MDa yielding an AAV particle with an assigned 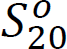 of 100 S. The three additional AAV species correspond to the same intact AAV shell, but are only partially filled, containing 90%, 80% and 70% by mass of the complete AAV DNA payload. The corresponding sedimentation coefficient for these three species was calculated from the difference between the sedimentation coefficient of the full, 100 S, and empty AAV, which was assigned a 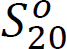 value of 63 S. The resulting 37 S difference was then multiplied by the percent mass of the full DNA payload (%DNA) present in a given partially filled AAV particle and added to the 63 S value for the empty protein shell as indicated using equation 1 to obtain the 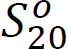 for partially filled AAV (note, this approach is analogous to Burnham et al.[7]):

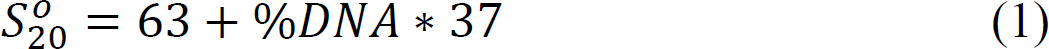

**Table 1.** Hypothetical AAV sample containing equal amounts of full and three partially filled AAV particles containing 90%, 80% and 70% of the full DNA payload.

| **\*%DNA** | **AAV particle MW MDa** | **Sed. coeff., $S_{20}^{\circ}$** | **$^{\S}$Sed. coeff., $S_{10}^{\circ}$** |
| --- | --- | --- | --- |
| **100** | **\*\*5.23** | **100** | **76.6** |
| **90** | **#5.10** | **##96.3** | **73.8** |
| **80** | **#4.98** | **##92.6** | **71.0** |
| **70** | **#4.83** | **##88.9** | **68.1** |

\* The % amount of the full DNA payload in the AAV particle
\*\* 3.90 MDa protein capsid MW + 1.33 MDa DNA MW

Sedimentation coefficients at 10°C, Table 1, were calculated using Equation 2:

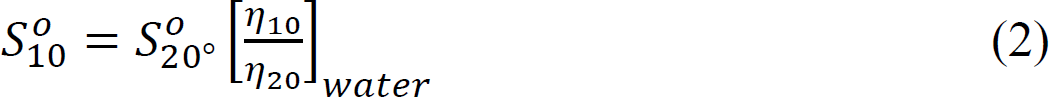

Changes in solution density and the partial specific volume of AAV in going from 20 °C to 10 °C were not considered significant in comparison to viscosity (η) changes.

### 2.2 Analytical Ultracentrifugation

All SV-AUC experiments were carried out in the Analytical Development department laboratories at Biogen located in Cambridge, Massachusetts. All AAV and solvent SV-AUC experiments were carried out on an Optima analytical ultracentrifuge equipped with both UV-Vis absorbance and refractometric (Rayleigh interferometer) detection. All runs used an An-60 Ti 4-hole rotor at 10 °C and 12 mm Epon double sector charcoal-filled centerpieces. AAV experiments were conducted at either15K rpm (standard SV-AUC), or 45K rpm (hs-SV-AUC), while solvent experiments were only conducted at 45K rpm (hs-SV-AUC).

#### 2.2.1 Cell filling

For hs-SV-AUC to function as intended, sample and reference sectors must be filled so that the radial difference between the reference solvent and sample menisci is ≤ 0.1 cm and the sample meniscus is at a radial position of about 6.1 cm or less. These filling specifications should be achieved easily in routine practice. Nevertheless, if these limits should be exceeded the resulting impact is very likely not that impactful, see Discussion section point “3”.

#### 2.2.2 Solvent mismatch SV-AUC experiments

Solvent mismatch SV-AUC experiments were performed for the 0.15 M NaCl solution, PBS buffer or AAV formulation buffer. All experiments were conducted at 45K rpm, at 10 °C. In all cases the sample sector solvent concentration was 5% lower than the reference sector concentration. Cell filling followed the protocol above with the sample meniscus at ∼6.1 cm and the reference meniscus at ∼6.0 cm (to achieve a menisci difference of about 0.1 cm between reference and sample sectors).

### 2.3 SV-AUC simulations

All SV-AUC simulations were produced using SEDFIT version 15.01b, employing the c(s) model for up to four non-interacting discrete (ideal behaving) species. All simulations resulted in 50 data scans computed using a radial resolution of 0.001 cm, and took into account rotor acceleration at 400 rpm/sec.

#### 2.3.1 AAV SV-AUC simulations

AAV SV-AUC simulations used the molecular weights (MWs) and temperature-corrected sedimentation coefficients shown in Table 1, an average partial specific volume, 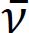, of 0.69 ml/g and a buffer density of 1.0 g/mL. AAV SV-AUC data for each temperature and rotor speed were generated using the time intervals indicated in Table 2 over a radial range of 6.1-7.2 cm, and with a signal to noise ratio (S/N) value of 10^3^ unless stated otherwise.

**Table 2.**
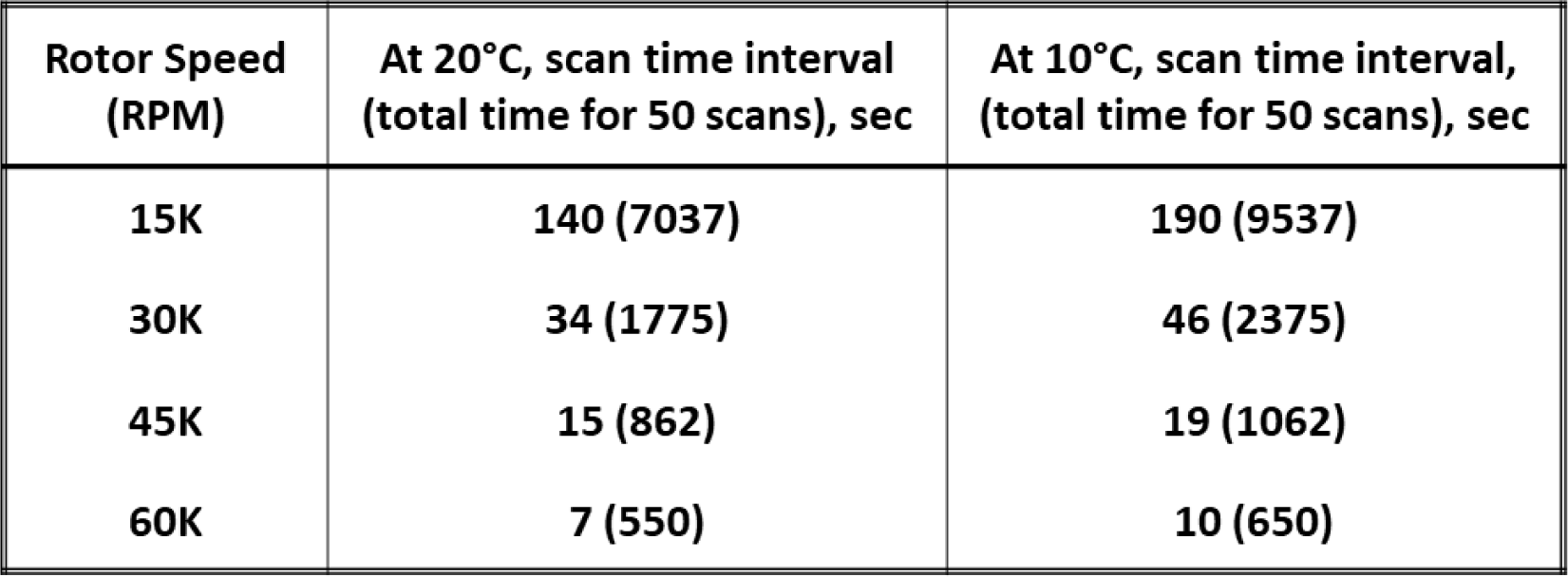
Required scan time interval and resulting total run time (), in seconds, to monitor the complete sedimentation of the full monomeric AAV particle in a total of 50 scans from 6.1-7 cm as a function of rotor speed and temperature.

#### 2.3.2 Simulating the solvent mismatch signal (in fringes) when doing SV-AUC with interference at different rotor speeds

The interference signal contribution arising from solvent mismatch was simulated for 0.15 M NaCl at the same rotor speeds and time intervals indicated in Table 2 for 10 °C using a 0.1 cm meniscus position mismatch and a 5% NaCl concentration mismatch (5% lower NaCl concentration in the sample sector). These specific levels of mismatch are unlikely to be exceeded by an AUC user and are considered *upper limit* values. Since the fringe displacement output of the interferometer is directly proportional to the concentration difference between the sample and reference sectors, our strategy was to simulate the sedimentation of the NaCl in each sector independently, then take the difference between the two sectors (see Figures 1S and 2S, which provides an outline and pictorial overview of the process, respectively). Specifically, solvent SV-AUC data was simulated separately for the reference sector between 6.0 – 7.2 cm at an initial concentration of 1.00 (arbitrary units) and the sample sector between 6.1 – 7.2 cm at an initial concentration of 0.95 (arbitrary units). For both simulations the S/N value was set to 10^6^, while the NaCl parameters were set as follows: MW= 58.45 Da, 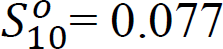 S (based on an average value of 0.1 S at 20 °C, from reported values [20,22,23]) and 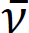 = 0.3 mL/g [24]. The resulting NaCl sedimentation scans for both sectors in arbitrary concentration units was then multiplied by the factor 25.5 (the concentration of 0.15 M NaCl in fringes) to convert them to fringe displacement readings (Figure 2SB). The conversion factor of 25.5 is based on a fringe count of 17 fringes at 670 nm obtained from a synthetic boundary experiment for 0.1 M NaCl in a 12 mm centerpiece [20]. For each time interval (scan) the expected solvent interference signal contribution, due to the differential NaCl sedimentation between the reference and sample sectors, was determined by taking the difference between the corresponding reference and sample signals (Figure 2SC). A “*solvent mismatch interference difference signal plot”* was generated by taking the difference between the two data scans corresponding to the first and last scan used in the c(s) analysis of the AAV hs-SV-AUC protocol (Figure 2SD).

**Figure 1.**
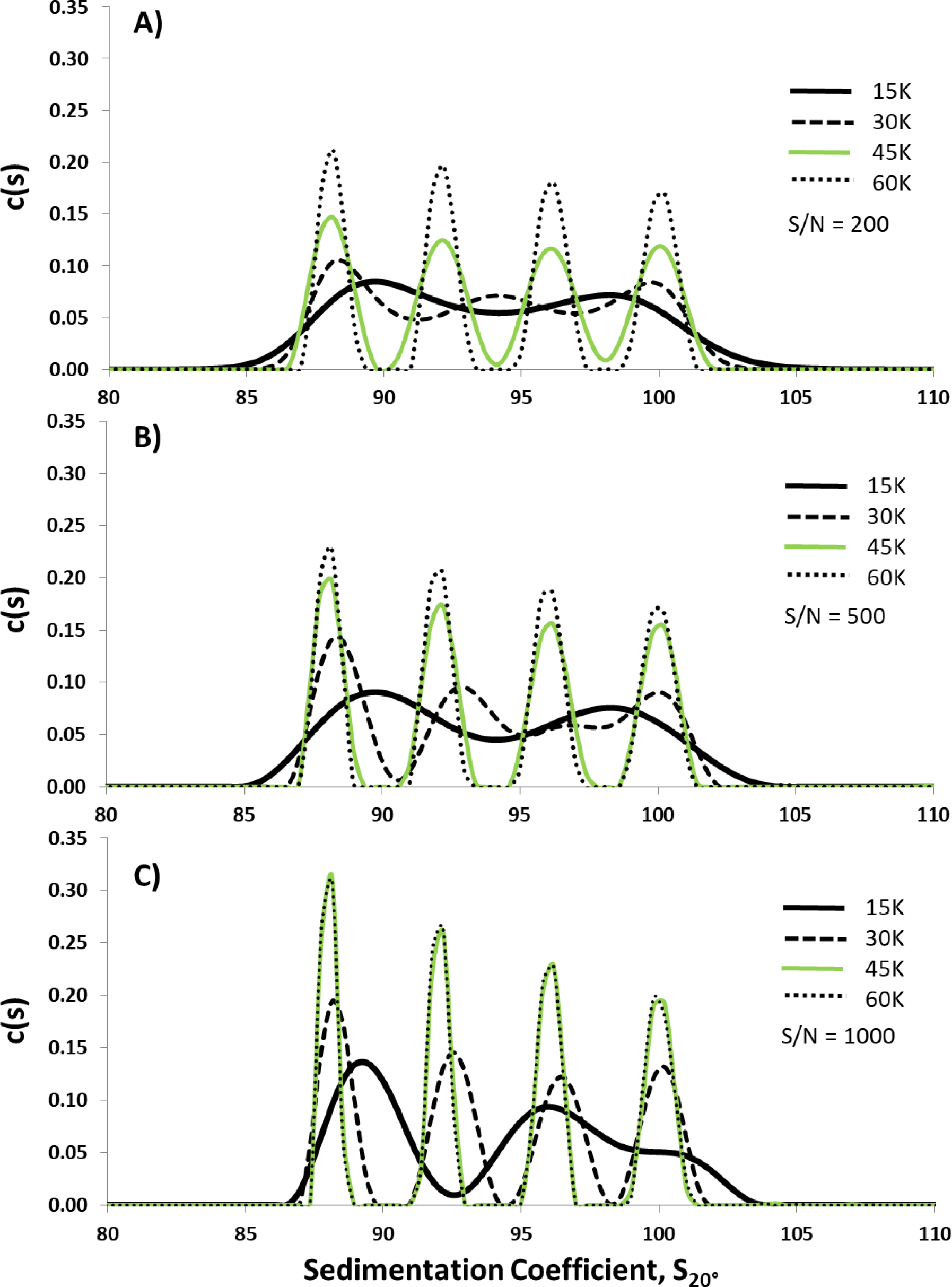
Computer SV-AUC simulation work on a hypothetical AAV sample (Table 1) in a 0.15M NaCl solution at 20 °C containing a 1:1:1:1 mix of AAV particles filled with different percentages (100%, 90%, 80% & 70%) of a viral vector DNA payload to assess the impact of rotor speed and S/N on the resolution of the resulting size distribution profile.

**Figure 2.**
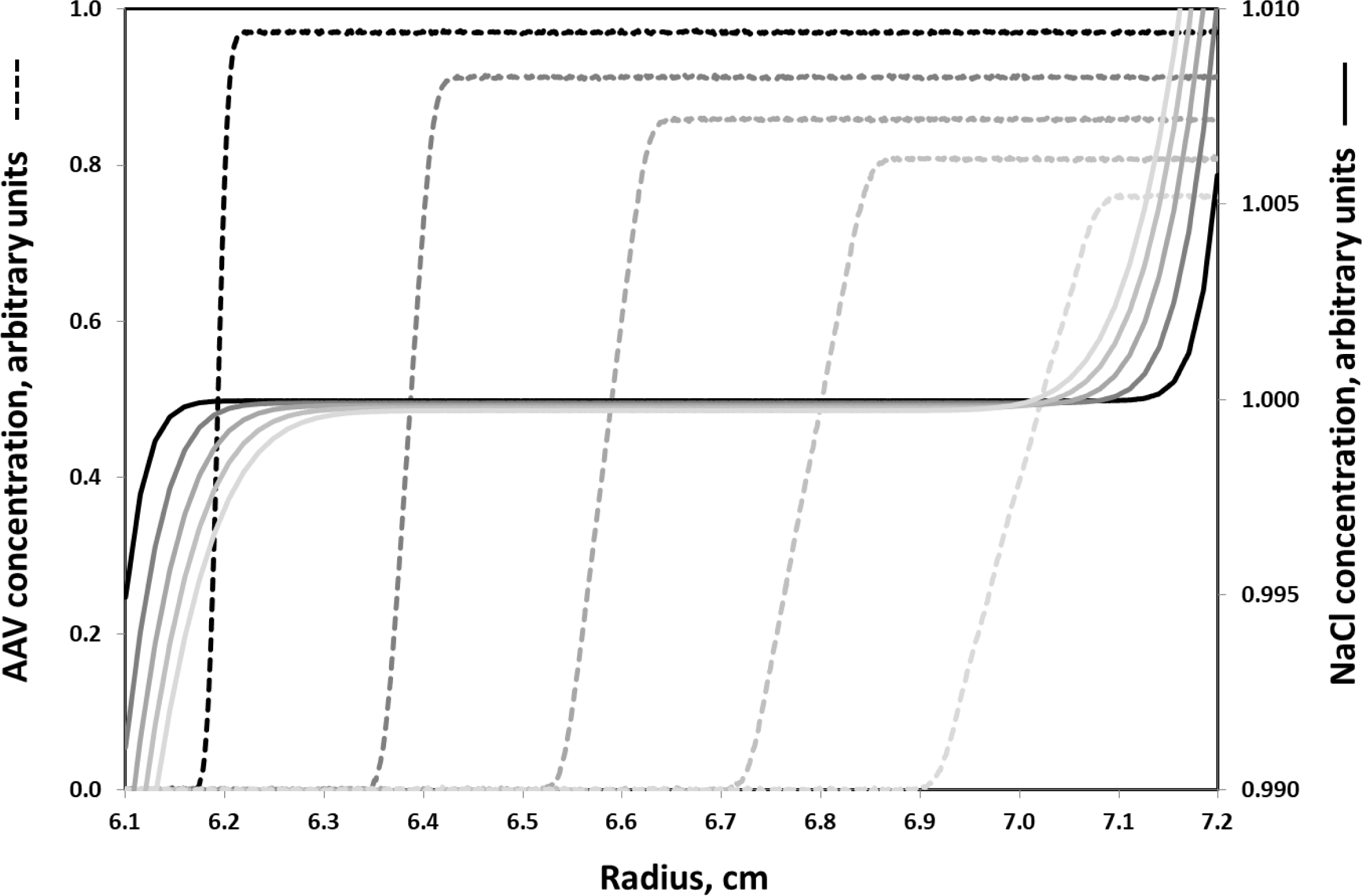
Overlay of the 1^st^, 10^th^, 20^th^, 30^th^ and 40^th^ data scans from a simulated hs-SV-AUC experiments on a hypothetical AAV sample (Table 1) in a 0.15 M NaCl solution, (dashed lines) and a 0.15 M NaCl solution (solid lines) run at 45K rpm at 20 °C using the same data scan time interval of 19 sec. In both cases the sample menisci are located at radius (r) of 6.1 cm with an arbitrary concentration (c) of 1.0 unit. Data scans shown in both cases, which were taken at the same time point have corresponding matching shades of black color (where black corresponds to the 1^st^ scan while the lightest shade of grey corresponds to the 40^th^ data scan). The resulting plot shows that the majority of time AAV migrates through a NaCl solution where there is little to no density (ρ) gradient (dρ/dr) stabilization (noting that dc/dr α dρ/dr). As a result, AAV material migrates through a large radial region (from about 6.3-7.0 cm) where it is exposed to the potential adverse effects of convection arising from the presence of very weak negative radial density gradients induced, for example, by the presence of increasing temperature with increasing radius.

### 2.4 Data analysis on AAV SV-AUC and hs-SV-AUC experiments and solvent mismatch hs-SV-AUC experiments

Data analysis on all AAV sedimentation velocity experiments used the c(s) distribution model in SEDFIT (version 15.01b), which assumes that all species migrate as discrete independent, ideal components. Low speed (15K rpm, 10 °C, SV-AUC) absorbance data were analyzed using a radial-time window of 6.15-7.15 cm and ∼9,500 sec. High speed (45K rpm, 10 °C, hs-SV-AUC) interference data used a radial-time window of 6.4-7.0 cm and ∼1,000 sec. All c(s) analyses employed data processing to remove both radial- and time-independent (RI and TI) noise signals. The steep boundaries encountered for AAV, especially for hs-SV-AUC, challenge finite element modeling [25,26]. Consequently both maximum entropy and 2^nd^ derivative (Tikhonov-Phillips) with Simplex regularization were used at a confidence level of 0.68. All c(s) vs S distribution plots presented here were obtained using 2^nd^ derivative regularization since it produced fewer erratic fits or extraneous peaks and avoided fusing closely spaced peaks [27]. Resolution was set at 250 and S_min_ and S_max_ values were set to 40 S and 130 S, respectively.

To determine the impact of solvent mismatch on AAV hs-SV-AUC data, solvent hs-SV-AUC data were analyzed by generating “*normalized solvent mismatch interference difference signal plot”* for each solvent by again taking the difference between the first and last scans in the same radial-time window used for AAV hs-SV-AUC experiments (i.e. from ∼6.4-7.0 cm and from the first scan until the scan acquired ∼1000 sec later). However, before subtraction, radial axial readings for each data scan were normalized by subtracting the middle radial reading (between 6.7-6.8 cm) for each scan from its corresponding radials readings.

## 3. Results

### 3.1 Effect of rotor speed and detector S/N on c(s) vs S distribution resolution

Generating high-resolution sedimentation coefficient size distribution plots is important for distinguishing the presence of unique components having closely-spaced sedimentation coefficients. AAV samples provide a good example where high-resolution analysis is important. In addition to the fully-intact AAV monomer, AAV samples often contain a variety of other particles, e.g., empty, partially filled, over filled virus and free DNA [13-16,28], that yield closely-spaced sedimentation coefficient profiles. Even homogeneous or heterogeneous aggregates (dimers) of some of these particles can form new particles that migrate close to the full monomeric AAV particle. To explore the impact of increasing rotor speed at 20 °C on resolution, SV-AUC data for a hypothetical AAV sample containing four-species (Table 1) were simulated using four different rotor speeds (15K, 30K, 45K and 60K rpm). For each rotor speed, the time interval between scans was chosen (Table 2) to generate 50 data scans [29] enabling the monitoring of the full AAV particle’s sedimentation from meniscus to cell base. Data were generated with stochastic noise levels consistent with the optimal S/N value for the detectors presently available: 200 for the XL-A/I absorbance detector, 500 for the Optima absorbance detector (at ∼1 OD signal) and 1000 for the interference detector (at ∼2-3 fringes) on either the XL-A/I or Optima. For each rotor speed, size distributions were assessed as described in Material and Methods. Figure 1 shows the increasing resolution achieved with increasing the rotor speed. While each increasing rotor speed improved the resolution, going from 45K rpm to 60K rpm offers only a marginal resolution gain while reducing sample throughput by requiring the use of the lower-capacity 4-hole (An-60 Ti) rotor. Resolution improved with higher S/N (Figure 1A-D), but the improvement is not as significant as that achieved by increased rotor speed. We conclude that 45K rpm is an optimal speed in terms of resolution and sample throughput.

### 3.2 Achieving convection-free sedimentation with hs-SV-AUC

Because AAV migration outruns the formation of a stabilizing solvent density gradient at 45K rpm and 20 °C (Figure 2), the hs-SV-AUC protocol is susceptible to sample convection due to thermal fluctuations [21], which can be of particular concern during the adiabatic cooling of the rotor on accelerating it to the running speed [21,30]. Even very small temperature-induced negative density gradients can cause convection due to the amplifying effect of the high gravitation field [30]. Lowering the rotor temperature from 20 °C to around 4°C, where 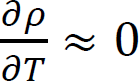, mitigates this problem. However, because rotor temperature equilibration can take a long time at 4 °C we investigated using 10°C for hs-SV-AUC, where 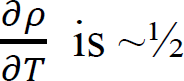 the value at 20 °C (see Figure 2S in reference [21]). Pre-equilibration of the rotor and samples to the lower temperature significantly reduces the equilibration time. However, to prevent sample leaks, cells should be re-torqued before running if they were loaded at room temperature.

### 3.3 Limitations of absorbance detection when doing hs-SV-AUC

To acquire 50 scans for analysis at 45K rpm requires that all cells are scanned every 19 sec or less (Table 2). Coupling this criteria with the scanning limit of the absorbance detector on the XL-A/I (30–60 sec/cell, depending on radial step size) and the Optima (7–20 sec/cell at a 10-micron step size) the best that could be achieved would be to acquire data from at most one cell (using only 1 wavelength) in the case of the Optima. Furthermore, even at the fastest scan rate there will be significant boundary movement during a scan. While it is possible to try to account for this data distortion, by assigning a unique time point to each radial reading during a given scan [31], the resulting increase in data analysis complexity exposes the analysis to opportunities for inaccuracies and added statistical uncertainties. Also, the sharp boundaries (or concentration gradient) encountered in hs-SV-AUC can result in schlieren distortion of the absorbance readings [32] and possible data analysis inaccuracies. Finally, the absorbance system optical resolution of both the XL-A/I or Optima is 50 μm [33], which can limit the system’s ability to follow sharp boundaries.

### 3.4 hs-SV-AUC using interferometry

The Rayleigh interferometer, a refractive detector, is capable of acquiring data from all samples in ∼5 sec, captures the entire cell image in a fraction of a second (i.e. no scanning), and has a radial resolution of ∼9 μm [18,34]. The detector also has comparable sensitivity to the absorbance detector (∼3.3 fringes/mg/ml versus ∼1 OD_280nm_/mg/ml or ∼5 OD_230_/mg/ml for proteins). The Optima provides fringe displacement measurements with an RMSD approaching ± 0.001 fringes (see Figure 3S), which is comparable to the absorbance detector precision at 230 nm. In addition to having a much wider dynamic range, the signal vs. concentration linearity of the interferometer is superior to the absorbance system, which is subject to stray light limitations at both low and high concentrations [18,33]. Its combination of rapid data acquisition, higher radial resolution, greater accuracy in tracing steep boundaries and superior S/N makes a properly focused interference system [35] the detector of choice for AAV hs-SV-AUC experiments.

**Figure 3.**
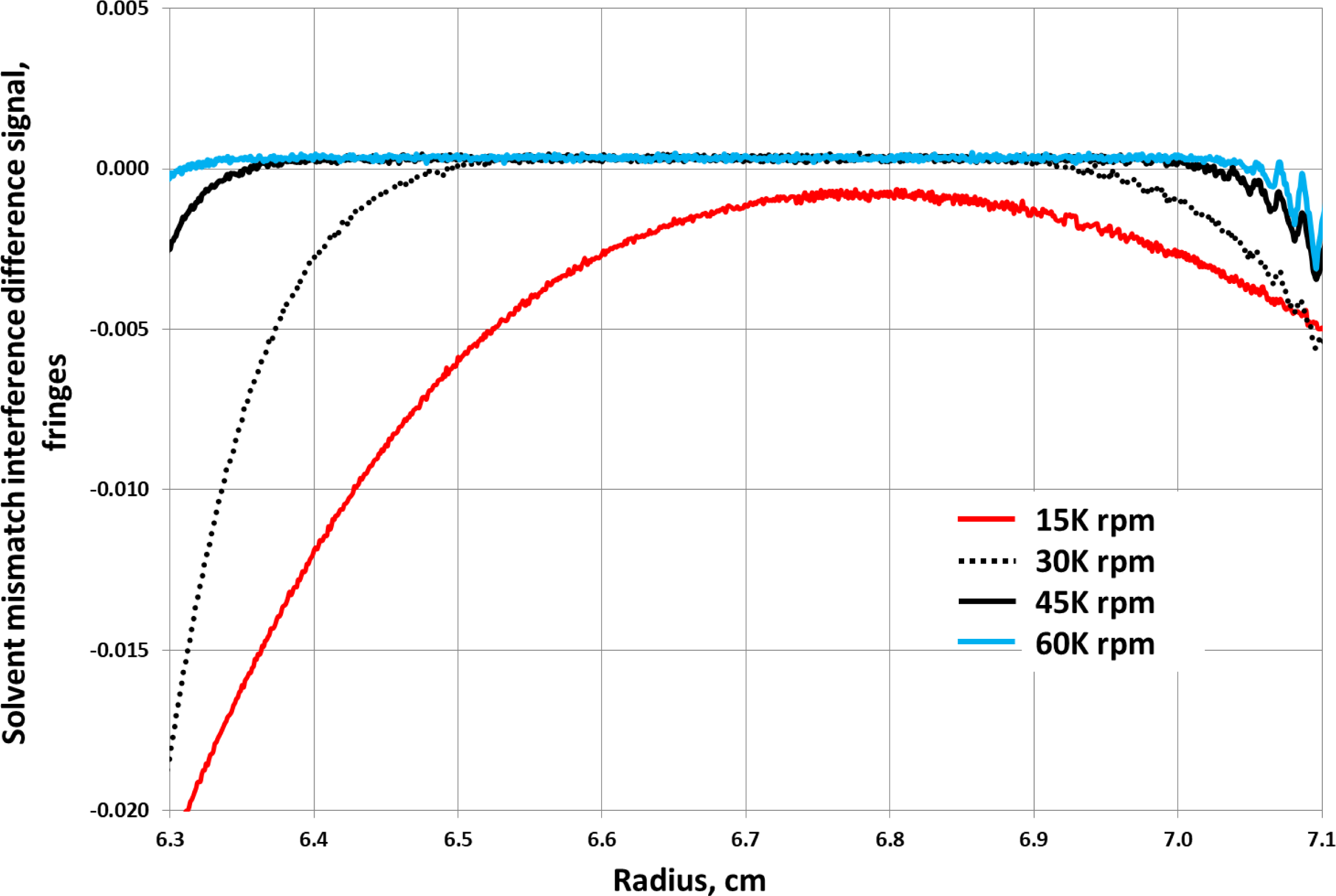
Overlay of the simulated “*solvent mismatch interference difference signal plots*” (see section 2.3.2) for 0.15 M NaCl (a proxy AAV formulation buffer or solvent) due to a 5% solvent compositional mismatch between the reference (0.15 M NaCl) and sample (0.1425 M NaCl) sectors of an AUC cell along with a 0.1 cm mismatch in menisci positions (due to difference in solvent filling volumes) between the reference (6.0 cm) and sample (6.1 cm) sectors of an AUC cell as a function of rotor speed at 10 °C: 15K (red), 30K (black, solid), 45K (black, dashed) and 60K (light blue) rpm.

However, the interference system is sensitive to all solution components, including the solvent salts and buffers and other formulation added excipients. Hence any difference in the solvent component sedimentation between the reference and sample sectors will give rise to refractive index changes that will contribute a signal to the total signal output from the interferometer. Even the slightest mismatch in the solvent component concentrations and difference in meniscus position between both sectors (due to slight difference in filling volume) will result in the superposition of the solvent component sedimentation with the macromolecular component signals of interest, distorting and complicating the data analysis. However, the short run time of hs-SV-AUC mitigates this problem by using a radial analysis region that is unperturbed by solvent boundaries. Only the flat signal decrease with time that is associated with radial dilution is present in the analysis region, and this contribution can be removed as RI noise by SEDFIT.

### 3.5 hs-SV-AUC overcomes the interference solvent mismatch problem

Solvent contributions to the interference signal were simulated for different rotor speeds with a 0.1 cm meniscus mismatch and 5% solvent concentration mismatch as described in Materials and Methods (and Figure 1S and Figure 2S) using 0.15 M NaCl as a proxy solvent. The resulting overlay of the “*solvent mismatch interference difference signal plot”* at each speed (Figure 3) indicate that AAV hs-SV-AUC experiments are unaffected by solvent sedimentation at 45K rpm and 10 °C, when limiting the radial-time analysis window to 6.4 – 7.0 cm and the first 1000 sec. Thus, the hs-SV-AUC protocol alleviates the need for extensive dialysis and meniscus matching. The question arises, though, whether the subset of data acquired using this radial-time window (Figure 4A) diminishes the analysis precision. Comparison of data analysis constrained to the protocol’s radial-time window (6.4-7.0 cm and 1000 sec) and analysis that encompasses the complete data range (6.15 – 7.15 cm and ∼1100 sec) shows minimal resolution reduction (Figure 4B) when analyzing simulated data using our hypothetical AAV test sample (Table 1). The simplicity (no dialysis or meniscus matching), increased resolution (Figure 1) and greatly shortened run time (< 20 min versus hours) makes AAV analysis by hs-SV-AUC protocol an attractive alternate approach to AAV SV-AUC conducted at much lower rotor speeds.

**Figure 4.**
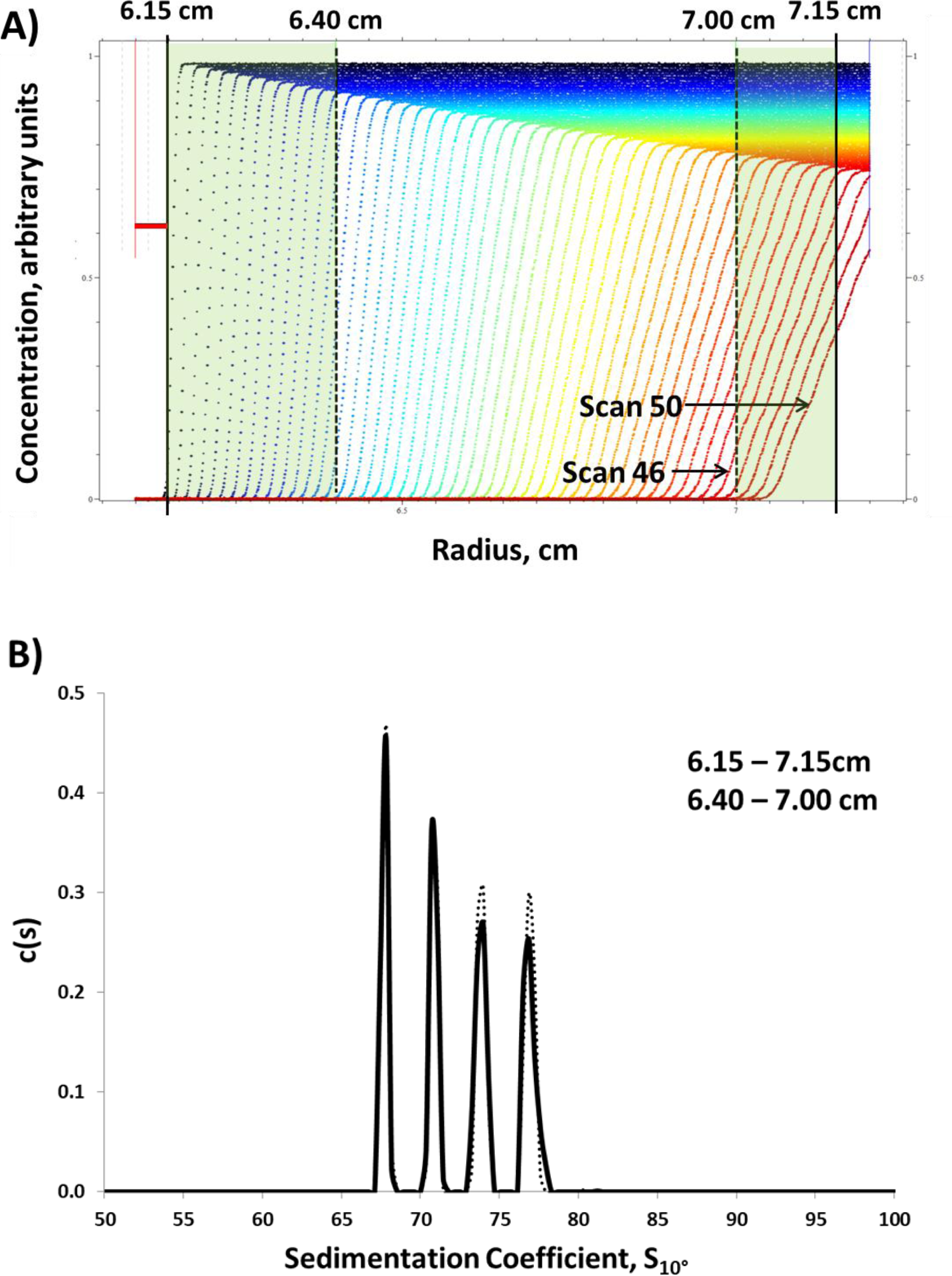
Comparison of the full (6.15-7.15 cm, ending with the 50^th^ data scan acquired at about 1100 sec from the start of the AAV hs-SV-AUC run) vs restricted (6.40-7.00 cm, ending with the 46^th^ data scan acquired at about 1000 sec from the start of the AAV hs-SV-AUC run) radial-time windows on the resulting computed AAV c(s) vs S distribution plot obtained from the same simulated AAV hs-SV-AUC data in 0.15 M NaCl at 45k rpm and 10 °C using a data scan time interval of 19 sec. **A)** Simulated 50 AAV hs-SV-AUV data scans used when doing AAV c(s) data analysis on the full radial-time window vs the restricted radial-time window where in the case of the latter, in terms of time only data scans 1-46 are used, while in terms of radius, data highlighted in light green are excluded. **B)** Resulting overlay of the computed AAV c(s) vs S distribution plots using the full radial-time window, dotted black line, vs the restricted radial-time window, solid black line. The horizontal light redline in “A” indicates sedimentation data lost due to the time delay needed for the rotor to reach running speed (for an acceleration rate of 400 rpm/sec, this amounts to about 120 sec for the An-60 Ti rotor).

### 3.6 Experimental verification of the solvent mismatch simulations

Experimental data acquired for three solvents using the hs-SV-AUC protocol and analyzed as described in Materials and Methods to generate “*normalized solvent mismatch interference difference signal plots”* are shown in Figure 5. For all three solvents, the meniscus and composition mismatches were similar to that used in the solvent (0.15 M NaCl) mismatch simulations (Materials and Methods). The resulting data are flat, in agreement with the simulations. Consequently, the interference signal contribution arising from solvent mismatch over the identified radial-time window of the AAV hs-SV-AUC protocol can be removed as RI noise in SEDFIT.

**Figure 5.**
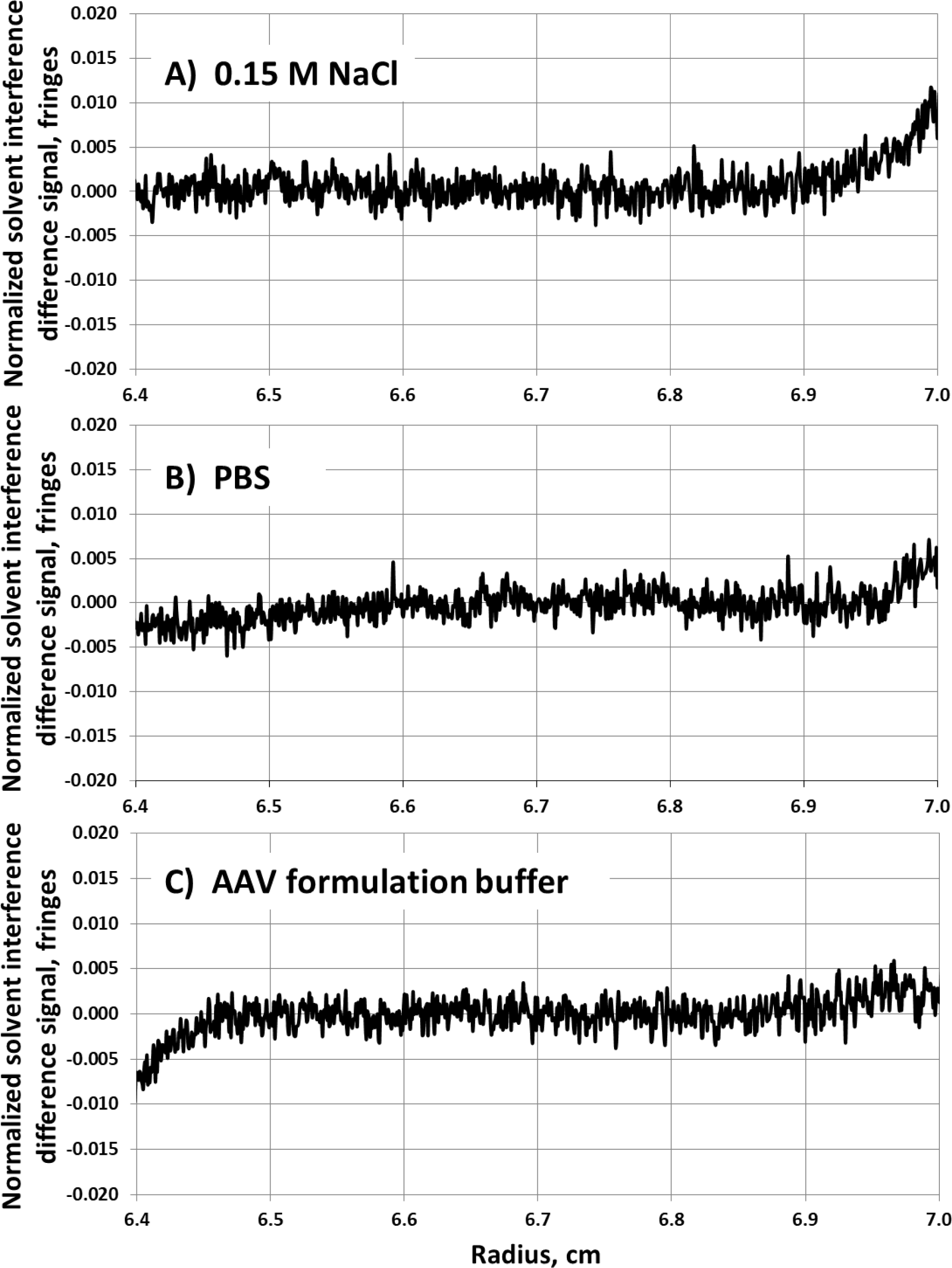
Experimental “*normalized solvent mismatch interference difference signal plot*” (see section 2.4) for three different solvents, generated using the AAV hs-SV-AUV protocol (45K rpm, 10 °C and the restricted radial-time window used in Figure 4). In all cases the cells were filled so the reference sector meniscus was at or near 6.0 cm, while the sample meniscus was at or near 6.1 cm (i.e. 0.1 cm mismatch), having a compositional mismatch of 5% (solvent composition was 5% lower in the sample sector than the reference sector): **A)** 0.15 M NaCl, **B)** PBS and **C)** AAV formulation buffer.

### 3.7 Experimental comparison of the same AAV sample run at 15K rpm vs 45K rpm (hs-SV-AUC) at 10 °C

Using a single AAV sample at two concentrations (0.2 and 0.45 fringes), three separate experimental runs were conducted using the AAV hs-SV-AUC protocol (45K rpm at 10°C) involving a total of 4 cells. Results from these 4 cells are shown in Figure 6 (see Figure 4S, for an overlay of these data). The same AAV sample was also run at 15K rpm and 10°C using absorbance and interference detection (at an AAV concentration of 2 OD_230nm_ ≈ 0.4 fringes involving a total of two cells. Results from these 2 cells, each generated from a separate SV-AUC run, are shown in Figures 7A and B reveal the reduced resolution in the size distribution relative to the data generated using the hs-SV-AUC protocol at 45K rpm at the same temperature. This difference in resolution, between the two the rotor speeds is more clearly demonstrated as an area normalized overlay plot shown in Figure 7C. The higher resolution afforded by hs-SV-AUC reveals the higher complexity of the distribution of partially filled capsids between the empty and filled AAV particles and resolves new peaks on either side of the full AAV particle. Both insights are unavailable from the low-speed data.

**Figure 6.**
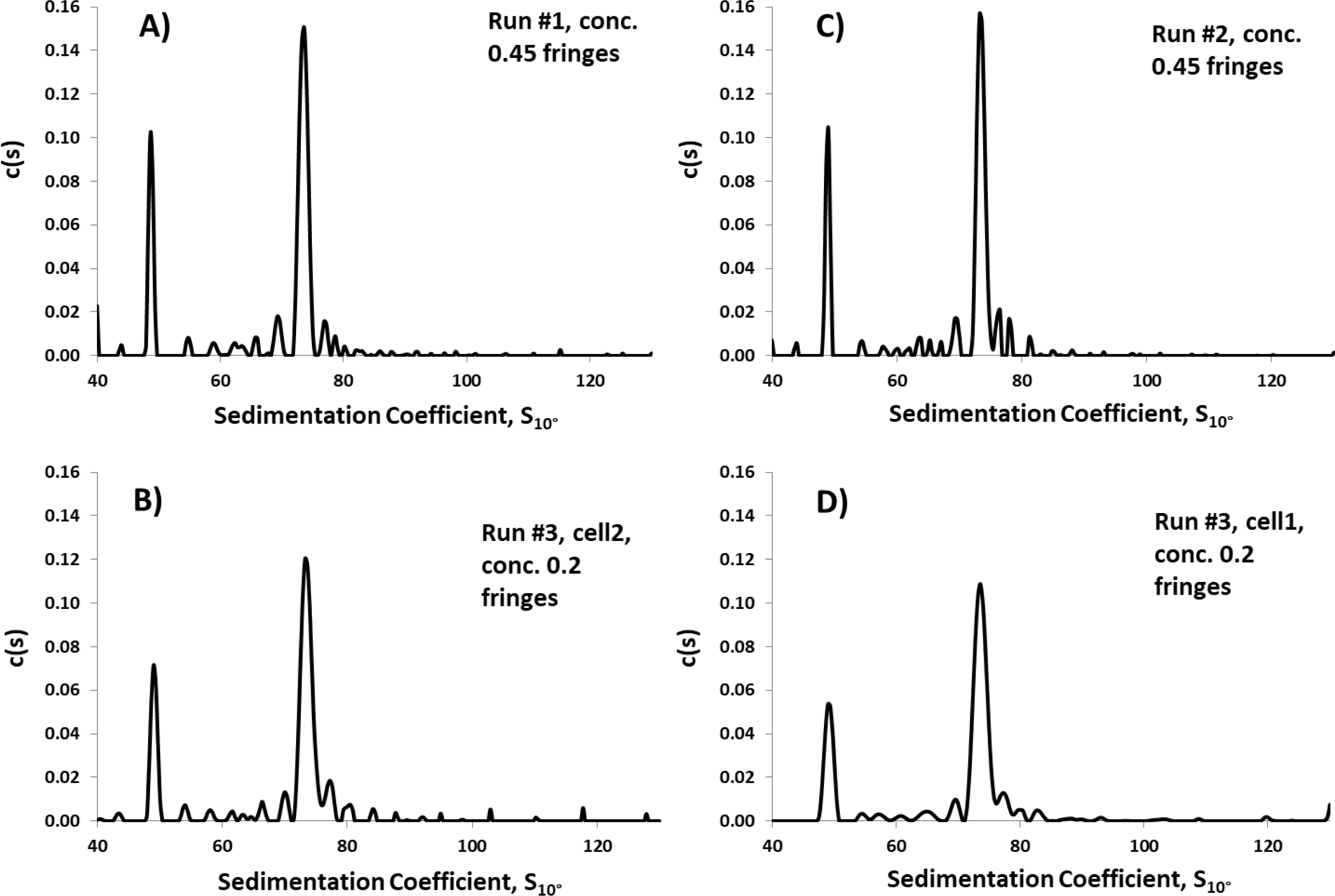
AAV c(s) vs S distribution plots obtained from three separate AAV hs-SV-AUC experimental run on different days on the same AAV sample (at concentrations indicated in units of fringes) at 45K rpm and 10 °C resulting in data from a total of 4 separate AUC cells **A-D**. It should be noted that plots **B-D** were normalized to the total fringe area in plot “**A”**. An overlay plot of these four area normalized AAV c(s) vs S distributions is shown in Figure 4S.

**Figure 7.**
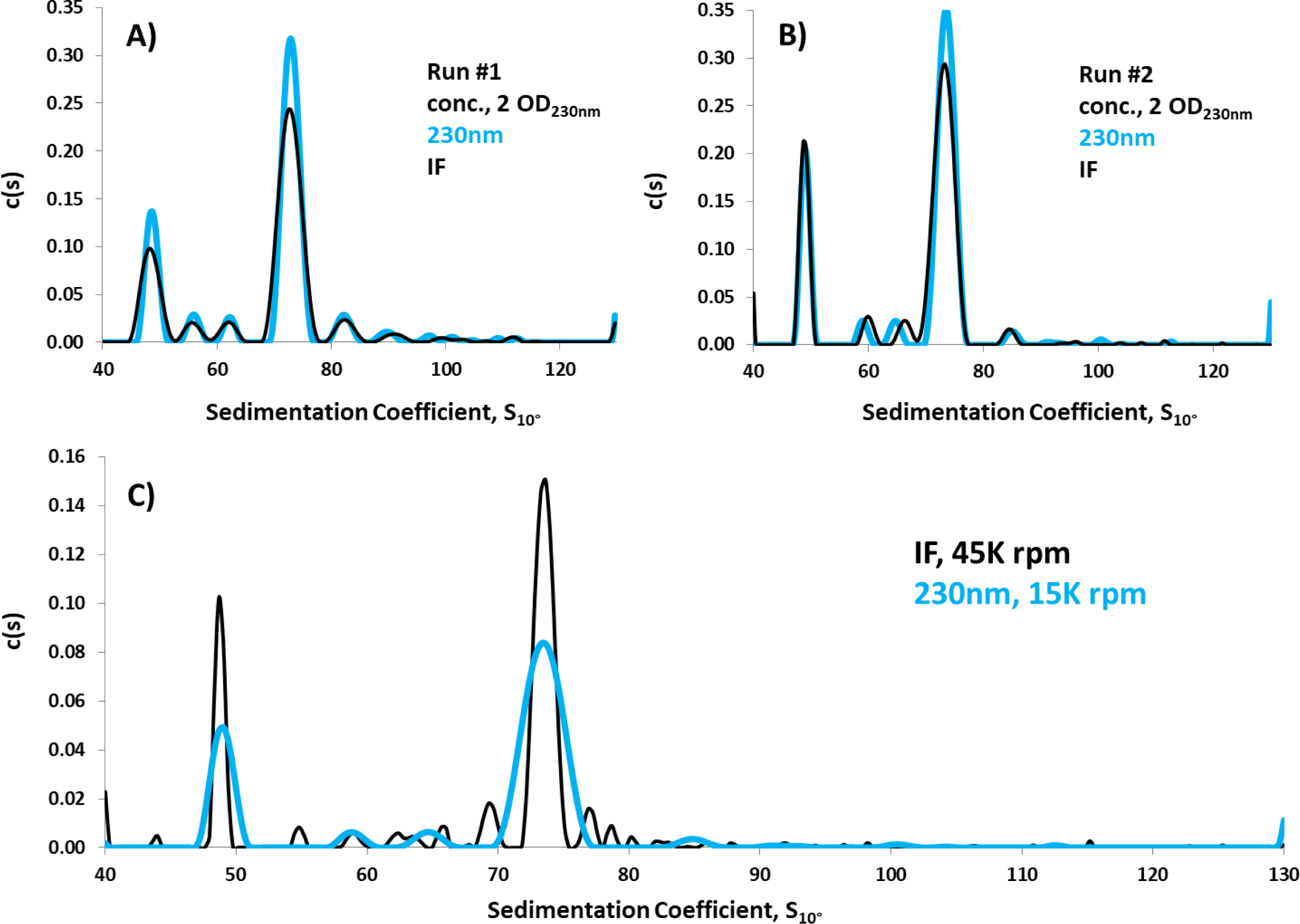
The overlay AAV c(s) vs S distribution plots in “**A**” and “**B**” were each obtained from two separate SV-AUC experimental runs conducted on different days at 15K rpm and 10 °C on the same AAV sample used in Figure 6 using absorbance, OD_230nm_, (light blue) and interference (black) detection. For each run the AAV c(s) vs S distribution plot from both detectors are provided as area normalized overlay plots (interference data was normalized to absorbance data). **C)** The resulting overlay of the AAV absorbance c(s) vs S distribution plot obtained at 15K rpm and 10 °C (light blue), shown in “**B**”, with the AAV c(s) vs S distribution plot at 45K rpm and 10 °C (black), shown in Figure 6A, for the same AAV sample (absorbance data was normalized to interference data).

A more quantitative comparison of data acquired with the same AAV sample analyzed at 15K and 45K rpm is provided in Table 3. Results show excellent agreement between the two sets of data for the sedimentation coefficients of the empty and full AAV particles, as well as the fractional amount of empty AAV particles. However, the percentages for partially filled, full, and aggregated (which may possibly include over filled) material showed noticeable differences. Some of this deviation could result from the different proportionality constants used to convert signal to concentration (extinction coefficient at 230 nm and refractive index increment at 670 nm) for the different chemical components of AAV (protein and DNA). However, given the proportionality constants for protein and DNA absorbance at 230 nm (extinction coefficient) and refractive index at 670 nm (refractive index increment) are similar [36], the enhanced resolution of material on either side of the full AAV particle peak at higher rotor speed (Figure 7C) is more likely the major contributing factor for the observed differences. Such data suggest that performing SV-AUC runs at lower speeds (i.e., at 15K rpm) might lead to an overestimation of the full particles fractional amount present in the AAV sample, when other AAV species are present in the sample with sedimentation coefficients that are fairly close to that of the full particle, and cannot be resolved at the lower speed.

**Table 3.** Quantitative analysis of the AAV sample run at 15K vs 45K rpm at 10 °C.

| Speed, detection | Sample ID | % Empty | Empty, S<sub>10°</sub> | % Partial | % Full | Full, S<sub>10°</sub> | % *Agg |
| --- | --- | --- | --- | --- | --- | --- | --- |
| Optima, 45K, IF | **RunId #1 | 20.6 | 48.7 | 11.6 | 58.0 | 73.6 | 8.3 |
| Optima, 45K, IF | **RunId #2 | 21.2 | 48.9 | 11.0 | 57.4 | 73.5 | 9.3 |
| Optima, 45K, IF | RunId #3_cell1 | 20.5 | 49.2 | 9.0 | 58.6 | 73.6 | 11.8 |
| Optima, 45K, IF | RunId #3_cell2 | 20.5 | 49.1 | 10.2 | 56.7 | 73.5 | 12.1 |
|  | Average | **20.7** | **49.0** | **10.4** | **57.7** | **73.5** | **10.4** |
|  | Std. Dev. | 0.3 | 0.2 | 1.1 | 0.8 | 0.1 | 1.8 |
|  | CV | **1.6%** | **0.4%** | **10.9%** | **1.4%** | **0.1%** | **18%** |
| Optima, 15K, Abs | **Run #1 | 20.1 | 48.7 | 7.8 | 61.0 | 72.8 | 11.5 |
| Optima, 15K, Abs | **Run #2 | 22.5 | 49.0 | 6.3 | 67.7 | 73.1 | 3.6 |
|  | Average | **21.3** | **48.8** | **7.0** | **64.3** | **73.0** | **7.6** |
|  | Std. Dev. | 1.6 | 0.3 | 1.1 | 4.7 | 0.3 | 5.6 |
|  | CV | **7.7%** | **0.5%** | **15.0%** | **7.3%** | **0.3%** | **74%** |
| Optima, 15K, IF | **Run #1 | 19.2 | 48.7 | 9.8 | 61.7 | 72.8 | 9.4 |
| Optima, 15K, IF | **Run #2 | 23.2 | 48.7 | 8.7 | 64.3 | 73.1 | 3.6 |
|  | Average | **21.2** | **48.7** | **9.2** | **63.0** | **73.0** | **6.5** |
|  | Std. Dev. | 2.8 | 0.0 | 0.8 | 1.8 | 0.2 | 4.1 |
|  | CV | **13.3%** | **0.0%** | **8.8%** | **2.9%** | **0.3%** | **63%** |
| Optima, 15K, Abs, IF | Average | **21.2** | **48.8** | **8.1** | **63.6** | **73.0** | **7.0** |
|  | Std. Dev. | 1.88 | 0.16 | 1.48 | 3.03 | 0.19 | 4.04 |
|  | CV | **8.9%** | **0.3%** | **18.3%** | **4.8%** | **0.3%** | **58%** |

\* Corresponds to AAV material migrating faster then the intact full AAV peak. As a result this may include monomeric overfilled AAV particles.
\*\* Sample concentration in this run is about twice that of samples in Run #3.

The results shown in Figures 6 and 7 clearly demonstrate the ability of hs-SV-AUC to acquire a more detailed characterization picture of the AAV heterogeneity and potentially a more accurate assessment of the active pharmaceutical ingredient, API, concentration. An example of AAV hs-SV-AUC data quality along with some statistical indicators of the quality of fit to experimental data can be found in Figure 3S.

## 4. Discussion

Robust methods for determining the size heterogeneity of the large (>30S), complex supramolecular biopharmaceuticals being developed for gene therapy (e.g. viral vectors and nanoparticles) are needed. When optimized, SV-AUC provides the most-accurate, broadest-range, and highest resolution size distribution analysis of any method [1–4]. Here, we describe the development of a high speed protocol, called hs-SV-AUC, that improves the size distribution analysis of supramolecular materials (in this case AAV viral vectors) that uses 45K rpm, low temperature (in this case 10°C) and interference optics. Both simulations and actual experimental AAV data reveal the following attributes about hs-SV-AUC:

1. *Improves particle size distribution resolution*. Since SV-AUC resolving power is proportional to rpm^2^, increasing the rotor speed used for AAV analysis from 15K rpm to 45K rpm improves component peak resolution. Higher resolution reveals important insights into drug product consistency, homogeneity and impurities, which are critical quality attributes (CQAs) in developing biopharmaceuticals. These CQAs may also shed light on the *in vivo* virus assembly process important in developing scalable economical production processes for the manufacturing of these drugs and in expanding our general knowledge concerning viral biological processes. Based on data we have generated we presently recommend doing AAV hs-SV-AUC at 45K rpm and 10 °C.
2. *Mandates the use interference (refractometric) detection for best results.* AAV component boundaries sediment quickly at 45K rpm making the absorbance detector scan time interval too long to be useful for hs-SV-AUC. Not only will there be too few scans for analysis, even for one sample, absorbance data will be potentially distorted by the resulting steep boundaries, the movement of the boundaries during the scanning process and the detector’s reduced spatial resolution. As a result, refractometric detection using the Rayleigh interferometer is required. Since the interferometer acquires the entire cell image in a fraction of a second, there is no discernable boundary movement during acquisition. Also, the entire data acquisition/storage process requires only a few seconds, allowing sufficiently short intervals between scans for acquisition using either the 4-hole or 8-hole rotor. Furthermore, the sensitivity of the interferometer, coupled with its excellent S/N characteristics and wide dynamic range, allows the analysis of dilute AAV solutions (a 1 OD_230nm_ sample in a 12 mm centerpiece provides a 0.2 fringe signal with a S/N of 200, Figure 3S). It is also worth noting that the interferometer may be used with solvents that are opaque at the wavelengths ordinarily used for AAV SV-AUC absorbance analysis (230 nm).
3. *Avoids the annoying challenges associated with solvent mismatch signal contributions to interference detection.* Interference detection is sensitive to the sedimentation of solvent components. Ordinarily, the need to properly match the reference and sample sector solvents requires exhaustive dialysis and meniscus-matching sample cells. However, for AAV and other supramolecular materials, confining data analysis to a radial region and run time window defined by 6.4 – 7.1 cm and the first 1000 sec results in a flat solvent plateau region through which the AAV boundaries are unaffected by solvent sedimentation. This feature enables SEDFIT to account for the solvent sedimentation as RI noise in computing the c(s) distributions. As a result, the hs-SV-AUC protocol avoids the need for exhaustive dialysis of AAV samples so long as the sample/reference sector meniscus position and formulation buffer compositional mismatches do not exceed ± 0.1 cm and ± 5%, respectively (see Figure 5S in the Supplemental Material concerning the robustness of the hs-SV-AUC method if these specific solvent mismatch limits are exceeded). Nonetheless, when first establishing an hs-SV-AUC method, after identifying the required sedimentation time needed to monitor the supramolecular structure of interest, it is recommended that a solvent mismatch experiment, as described in section 2.2.2 and shown in Figure 5, be conducted so that the appropriate radial-time window can be determined for the specific solvent.
4. *Uses a low temperature (10 °C):* Using a lower temperature for AAV analysis has two benefits: 1) the higher viscosity of aqueous solutions at 10 °C versus 20 °C slows sedimentation such that *S*_10_° = 0.766 · *S*_20_°. This slower sedimentation allows for more useful scans to be acquired at higher rotor speeds, and 2) the lower temperature reduces the chance for thermally-induced convection. A temperature of 10 °C seems to be adequate, but lower temperatures are even less prone to convection [21]. Thermally-induced convection is most frequently encountered during or shortly after the acceleration phase of the rotor to its running speed. Here adiabatic cooling resulting from rotor expansion causes the temperature control system to call for heat. It should also be noted that during acceleration, compression by the cells at the outer edge of the rotor holes leads to localized rotor warming that is not sensed by the temperature control system. As a result, the extent and rate of temperature change is controlled by the acceleration rate and the final speed of the rotor. Both the XL-I and Optima have a maximum acceleration rate of 400 rpm/sec (which for the former is the default setting). As a result, a slower rotor acceleration rate (200 rpm/sec or possibly even 100 rpm/sec) might allow sufficient heat flow within the rotor to balance the temperature changes occurring during acceleration, thus minimizing the extent and rate of temperature change reducing the chance of convection. For the XL-I user such an approach is doable since the user has some level of control over rotor acceleration, while in the case of the Optima this capability is not available to the Optima user (however, multi-speed SV-AUC [37,38] might offer an alternative to mimic acceleration control). The tradeoff, of course, is that it will take longer to reach speed, thus lengthening a run, which could lead to a narrowing of the radial-time window.
5. *Improves sample throughput.* Due to the high rotor speed employed with hs-SV-AUC, run times are short, typically < 20 min. at 10 °C. This allows for two or three hs-SV-AUC runs to be conducted in single day, with each run carrying three/four (4-hole rotor) or seven/eight (8-hole rotor) cells depending on the instrument being used, Optima/XL-I. This sample throughput is significantly greater than that offered by a typical SV-AUC low-speed protocol. Having a second set of AUC cells available that can be cleaned, filled and stored in a refrigerator facilitates higher throughput. Having the replacement cells at or just below the running temperature will avoid problems in inserting and aligning the cells in the rotor hole. To prevent cell leaking, just prior to putting the cells in the rotor be sure to re-torque them. Once the rotor is removed from the centrifuge, the cell replacement process should be conducted as quickly as possible to minimize rotor temperature change and water condensation on the rotor (remove any condensation by wiping with a lint-free cloth before placing the rotor into the rotor chamber). By minimizing temperature changes and condensation, the time needed for pump-down and thermal equilibration will be minimized. Once the rotor has reached the run temperature, allow ∼2 hours to assure thermal equilibrium before starting the next run. During this interval, cells from the first run can be prepared for the next run.

Although we have focused on the application of hs-SV-AUC to AAV characterization, the hs-SV-AUC protocol can be applied to a wide range of other gene therapy products (e.g., other viral vectors, virus-like particles (VLPs) and nanoparticles) of similar or greater size and sedimentation coefficient values. Likewise, hs-SV-AUC should find useful applications in polymer and material science where a broader range of large more complex supramolecular structures also exists.

Nevertheless, in developing a hs-SV-AUC protocol for other biological or non-biological systems there are considerations to be aware of, particularly concerning the radial-time window used. There are also differences to consider when operating the Optima’s interferometer relative to the more common XL-I’s interferometer.

For a given rotor speed the radial-time window associated with hs-SV-AUC puts limits on the upper and lower range of S values that can be characterized. In the case of AAV the upper and lower size limit was approximated using hs-SV-AUC simulations using a 2^nd^ hypothetical AAV (consisting of equal amounts of intact full AAV monomer and several aggregates of this monomer, of specific size and configuration) and a hypothetical monoclonal antibody, mAb, sample (consisting of equal amounts of the monomeric mAb and several aggregates of this monomeric mAb, of specific size and configuration), respectively. In the case of the upper S limit results showed that aggregates with S values greater than about 150 S_10°_ will migrate through the radial-time window before data acquisition starts (Figures 6SA & B). In the case of the lower S limit material having S values less than about 20 S10° will not enter the radial-time window by the time data acquisition has terminated (see Figures 6SC & D). In general, using the hs-SV-AUC protocol for AAV, samples that contain material having S values *outside* the range of about 20-150 S_10°_ will not be detected. Therefore, it is recommended that an initial standard 2-3 hour, lower-speed (15K rpm) run be conducted to ensure that there is no material > 150 S_10°_ or < 20 S_10°_ before 45K rpm is used for routine analysis. If there is material outside this range, opportunities for extending the limits for the AAV hs-SV-AUC do exist, see the discussion below.

An approach to increase the upper S limit when doing AAV hs-SV-AUC, in order to characterize the presence of larger AAV material (aggregates), shortening the data acquisition interval from 19 sec to ∼5 sec (the limit for the interferometer on both the Optima and XL-I) increases the number of data scans collected. In the case of conducting AAV hs-SV-AUC using the XL-I this increase in scan rate at 45K rpm at 10 °C increases the upper S limit value to ∼300 S_10°_ using the same radial-time window (see Figure 7S). However, it should be noted that the ability to implement this approach on the Optima is not available due to the automated setup operation of its interferometer, see the brief discussion in the last paragraph of this section.

Two additional strategies to expand the upper S limit value for AAV hs-SV-AUC is the following:

1) Slow down the sedimentation process by reducing the temperature from 10 to 5 °C. Reducing the temperature not only allows capturing the sedimentation of the larger species, but also further reduce the potential for encountering convection and, from preliminary simulation solvent mismatch hs-SV-AUC work, may even slightly expand the radial-time window.

2) Perform a multi-speed hs-SV-AUC [37,38] experiment in which the rotor is set to collect data at different speeds during different stages of the same SV-AUC run. However, simulation and actual solvent hs-SV-AUC experiments should be carried out to assess the impact of solvent mismatch contribution to the final interference signal at each of the rotor speeds.

An approach to decrease the lower S limit when doing AAV hs-SV-AUC, in order to characterize For the presence of smaller AAV material (e.g., AAV protein, DNA, and capsid and DNA fragments), < 20 S_10°_, the unique independent operation of the Optima’s interferometer and absorbance detectors allows much lower S values to be monitored. In this scenario, high-S material is monitored using the interference data, adhering to the hs-SV-AUC radial-time window, while the low-S material is monitored using a much wider radial-time range of a SV-AUC run with absorbance detection. In this situation the absorbance system limitations described above are not an issue for the slower moving material. This dual-detector arrangement expands the lower S range down to about 1 S_10°_. It does so, however, by increasing the run time from ∼20 minutes to 2 hours or more. Nevertheless, dual-detector analysis on the Optima can expand the normal S range covered in an AAV hs-SV-AUC experiment from about 1-150 S_10°_. Increasing the AAV loading concentration improve hs-SV-AUC S/N, and facilitates the sensitivity in detection of smaller (S_10_ < 20) and lower concentration AAV contaminants, surpassing the capabilities of low speed SV-AUC using absorbance detection.

The Optima analytical ultracentrifuge manufacturer chose to simplify the operation of the interferometer on the Optima by automating the task of setting the light source timing (typically done on the XL-I by conducting a separate SV-AUC run at the same target rotor speed). While the simplification is appreciated, *every time* the Optima interference detector is used, a laser setup procedure is conducted before data acquisition starts. Unfortunately, the time required to perform this task (which varies somewhat from run to run), after the rotor reaches run speed leads to a loss of useful data acquisition time. In the case of AAV hs-SV-AUC at 45K rpm at 10 °C this loss of data is fortunately not significant, see Figures 8SB-D. Nevertheless, the ability to extend the upper S value when doing AAV hs-SV-AUC at 45K rpm, as discussed above, is limited significantly for the Optima (Figure 6SB). Optima users need to be aware of this attribute since this delay can be a point of concern when applying hs-SV-AUC to other biopharmaceutical systems. Ideally, the ability to perform the laser setup procedure independently from the run itself, or at a minimum, the ability to request that the instrument skips this setup at the beginning of every run are two strategies that can overcome this problem.

## 5. Conclusion

The size distribution of a biopharmaceutical is an important CQA for assessing the purity and the manufacturing consistency of these drugs, which translates into their therapeutic dosing accuracy. Although other biophysical techniques can provide size distribution information on AAV, and other viruses and nanoparticles [17,39,40], and do so in many cases with higher sample throughput and/or requiring very small amounts of sample material, when properly optimized SV-AUC provides the broadest, most accurate and most detailed size distribution on a AAV sample in its *native state* of any method [1–4]. In this paper, we show that for AAV, hs-SV-AUC can further improve this capability of SV-AUC to provide even higher resolution size distribution information and allows for a higher throughput by eliminating sample preparation steps (such as dialysis) and minimizing the experiment run time (< 20 min). While the experimental parameters determined in this manuscript for the optimized hs-SV-AUC experiment (45K, 10 °C, interference detection, etc…) can be used for the analysis of most AAV samples, the numbers are not meant to be prescriptive. Rather, the authors aim to provide AUC users with the strategies required to develop and optimize their own hs-SV-AUC method, or at a minimum, verify that the method parameters determined here are suitable for their buffer system (mismatch issue). Other variations of the hs-SV-AUC experiment are possible. For instance, when high throughput is not required (in a QC lab, for instance) one can imagine that using absorbance detection at a slightly lower speed (40K rpm) overcomes the challenges associated with the steep boundaries. Alternatively, dropping the temperature below 10 °C enables further slowing down of sedimentation of larger species and can potentially capture large AAV aggregates often present in in-process intermediate samples. Regardless of the intended use, hs-SV-AUC offers multiple advantages over the classical SV-AUC experiment and should become the preferred method for the analysis of AAV samples.

