## Supplemental Figures 1S-8S for "Rapid high-resolution size distribution analysis for adeno-associated virus using high speed SV-AUC"

**Figure 1S**

**Supplemental Material**

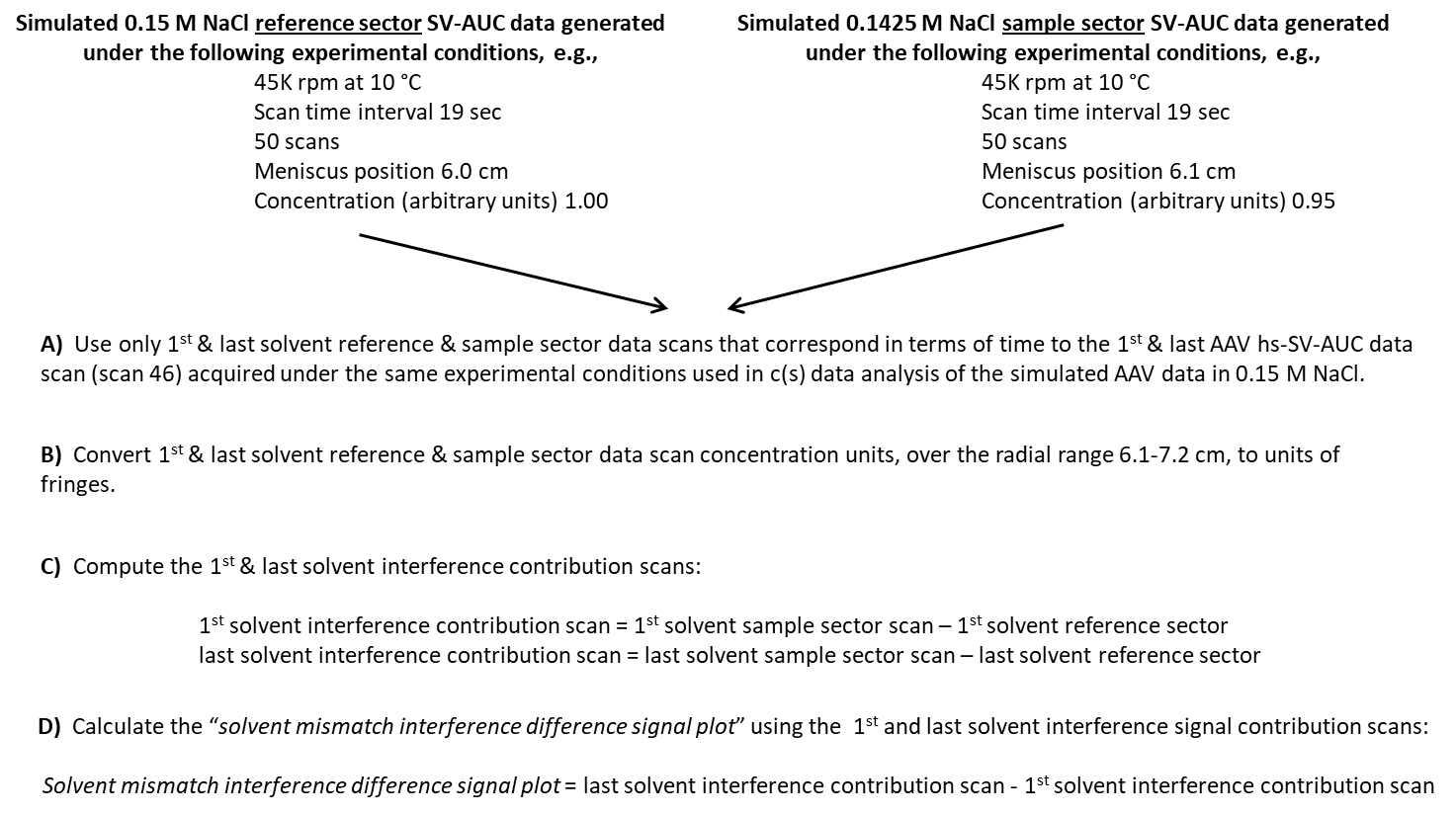

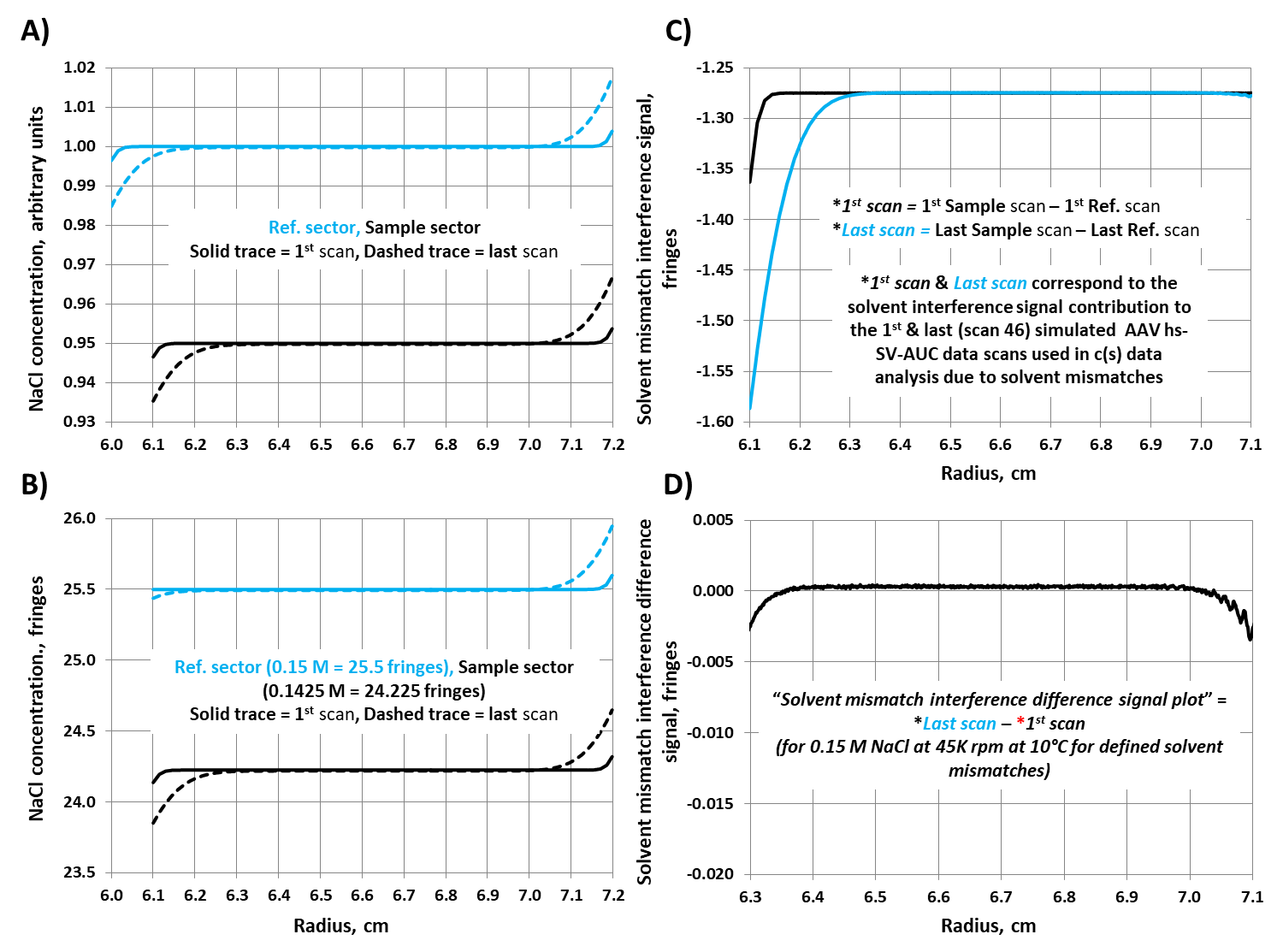

**Figure 2S**

**Figure 3S**

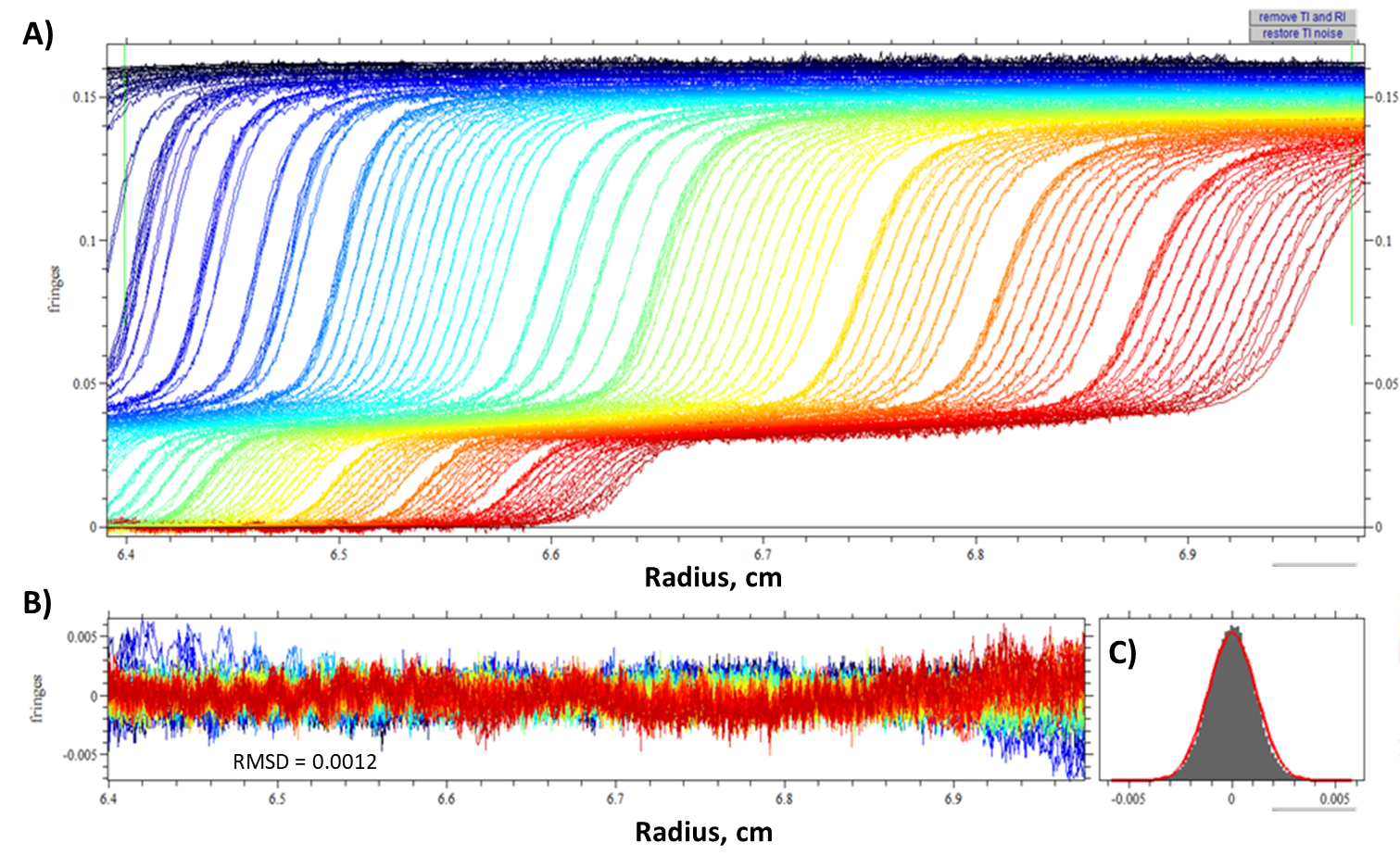

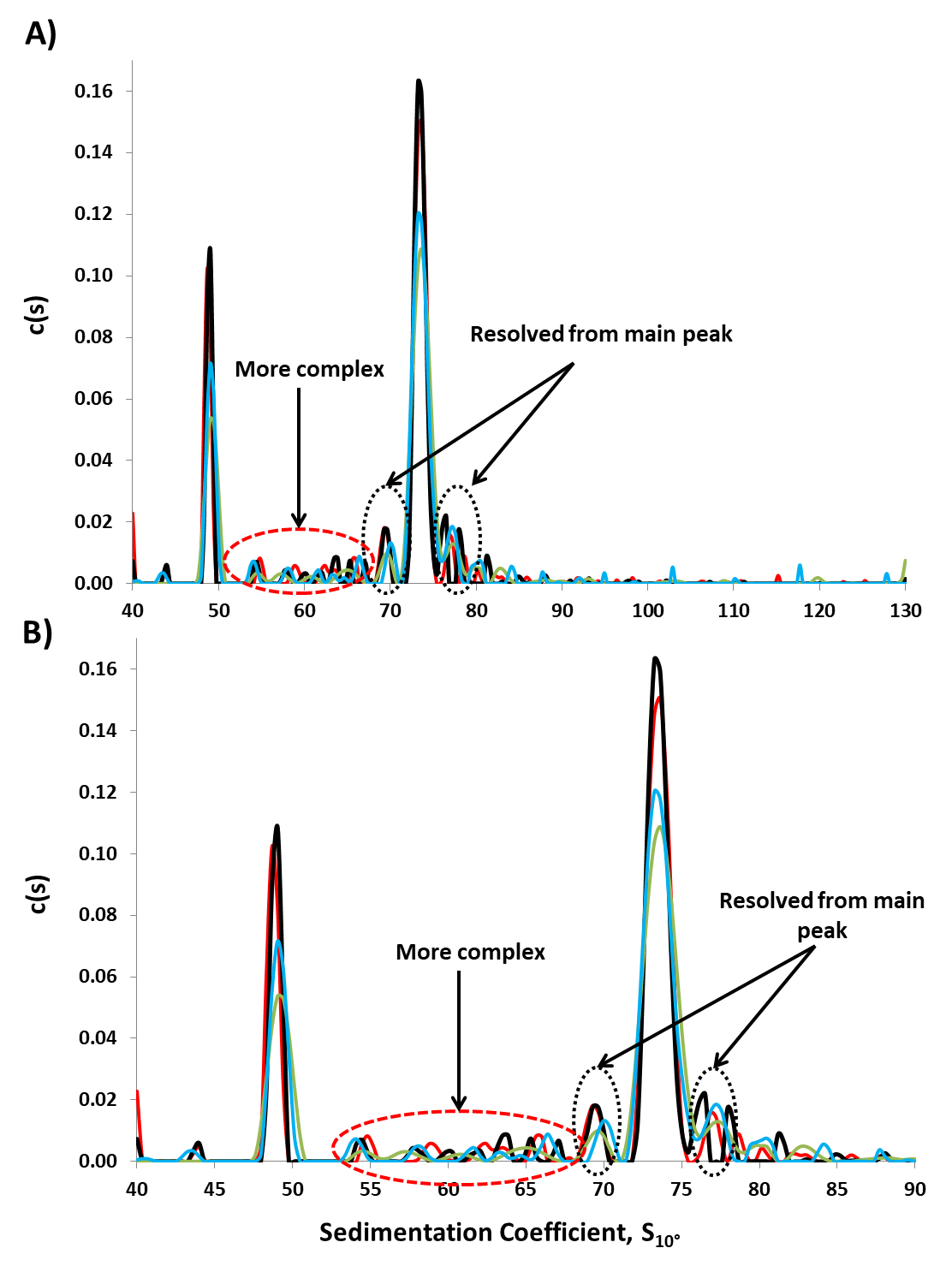

**Figure 4S**

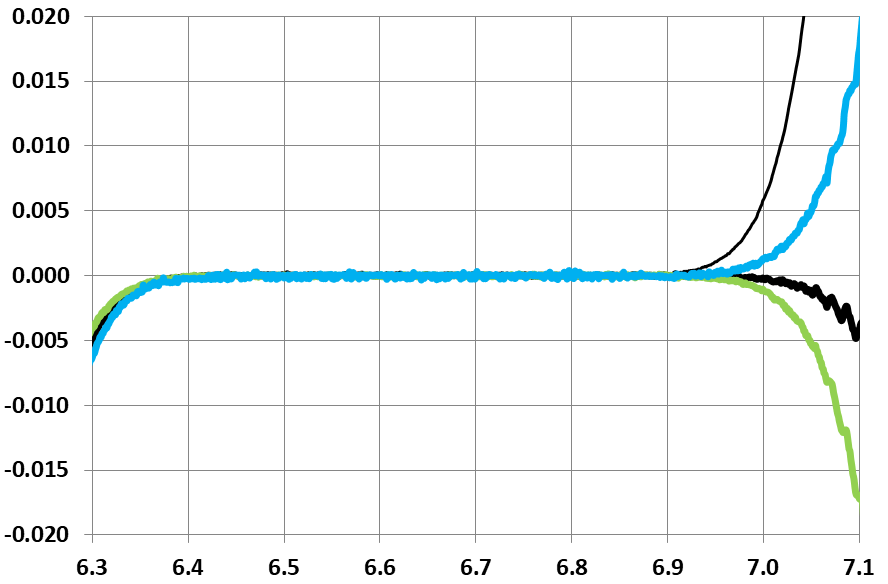

**Ref: 6.0, 0.15 M NaCl, Sample: 6.1, 0.1425 M NaCl:** 0.1 cm, -5%

**Ref: 5.9, 0.15 M NaCl, Sample: 6.1, 0.1425 M NaCl: 0.2 cm, -5%**

Ref: 6.0, water, Sample: 6.1, 0.1425 M NaCl: 0.1 cm, +100%

**Ref: 6.0, 0.15 M NaCl, Sample: 6.1, 0.18 M NaCl: 0.1 cm, +20%**

**Radius, cm**

**Solvent mismatch interference difference signal, fringes**

**A)**

**Figure 5S**

|  |  |  |  |  |
| --- | --- | --- | --- | --- |
| 6.1 | -0.324868 | -0.39687 | -0.182004 | -0.309216 |
| 6.101 | -0.321247 | -0.391846 | -0.182564 | -0.309113 |
| 6.102 | -0.317575 | -0.386721 | -0.183075 | -0.308986 |
| 6.103 | -0.313928 | -0.381672 | -0.183635 | -0.309368 |
| 6.104 | -0.310332 | -0.376623 | -0.184094 | -0.309496 |
| 6.105 | -0.306558 | -0.371498 | -0.184681 | -0.309369 |
| 6.106 | -0.302988 | -0.366474 | -0.185242 | -0.309522 |
| 6.107 | -0.299291 | -0.3614 | -0.185854 | -0.309726 |
| 6.108 | -0.295619 | -0.356299 | -0.186186 | -0.310032 |
| 6.109 | -0.292049 | -0.351302 | -0.186798 | -0.309929 |
| 6.11 | -0.28825 | -0.346151 | -0.18746 | -0.309853 |
| 6.111 | -0.284705 | -0.341127 | -0.187945 | -0.309955 |
| 6.112 | -0.280957 | -0.336027 | -0.188481 | -0.310108 |
| 6.113 | -0.27731 | -0.330952 | -0.18899 | -0.310236 |
| 6.114 | -0.273536 | -0.325904 | -0.189347 | -0.310159 |
| 6.115 | -0.26966 | -0.320957 | -0.189373 | -0.309675 |
| 6.116 | -0.266192 | -0.316545 | -0.188149 | -0.307456 |
| 6.117 | -0.262851 | -0.312133 | -0.186849 | -0.304829 |
| 6.118 | -0.259486 | -0.307748 | -0.185701 | -0.302586 |
| 6.119 | -0.25617 | -0.303387 | -0.184426 | -0.300061 |
| 6.12 | -0.252805 | -0.299001 | -0.183151 | -0.297766 |
| 6.121 | -0.24949 | -0.29459 | -0.181902 | -0.295547 |
| 6.122 | -0.246098 | -0.290229 | -0.180805 | -0.293304 |
| 6.123 | -0.242834 | -0.285843 | -0.179505 | -0.290957 |
| 6.124 | -0.239494 | -0.281482 | -0.178306 | -0.288484 |
| 6.125 | -0.236102 | -0.277071 | -0.177107 | -0.286215 |
| 6.126 | -0.232761 | -0.272685 | -0.175832 | -0.283613 |
| 6.127 | -0.229472 | -0.268324 | -0.174609 | -0.281344 |
| 6.128 | -0.226132 | -0.263938 | -0.173359 | -0.279126 |
| 6.129 | -0.222816 | -0.259578 | -0.172212 | -0.276575 |
| 6.13 | -0.219527 | -0.255319 | -0.170809 | -0.274051 |
| 6.131 | -0.216543 | -0.251622 | -0.16882 | -0.270915 |
| 6.132 | -0.213509 | -0.247848 | -0.166703 | -0.267268 |
| 6.133 | -0.210551 | -0.24415 | -0.164944 | -0.264234 |
| 6.134 | -0.207593 | -0.240427 | -0.162981 | -0.260664 |
| 6.135 | -0.204635 | -0.23673 | -0.16094 | -0.257553 |
| 6.136 | -0.201601 | -0.232956 | -0.159028 | -0.254492 |
| 6.137 | -0.198617 | -0.229259 | -0.157116 | -0.250872 |
| 6.138 | -0.195659 | -0.225561 | -0.155229 | -0.247837 |
| 6.139 | -0.192727 | -0.221863 | -0.153239 | -0.244216 |
| 6.14 | -0.189641 | -0.218064 | -0.151327 | -0.241182 |
| 6.141 | -0.186683 | -0.214366 | -0.149261 | -0.237561 |
| 6.142 | -0.183725 | -0.210618 | -0.147298 | -0.234449 |
| 6.143 | -0.180792 | -0.206946 | -0.14536 | -0.231134 |
| 6.144 | -0.177809 | -0.203198 | -0.143524 | -0.227871 |
| 6.145 | -0.17508 | -0.199704 | -0.14156 | -0.224708 |
| 6.146 | -0.172683 | -0.196568 | -0.139852 | -0.221929 |
| 6.147 | -0.170133 | -0.193507 | -0.137838 | -0.218334 |
| 6.148 | -0.167558 | -0.190396 | -0.1359 | -0.215325 |
| 6.149 | -0.164931 | -0.18726 | -0.133834 | -0.211958 |
| 6.15 | -0.162407 | -0.1842 | -0.131896 | -0.208694 |
| 6.151 | -0.159755 | -0.181089 | -0.129856 | -0.205634 |
| 6.152 | -0.157154 | -0.177978 | -0.127969 | -0.202141 |
| 6.153 | -0.154706 | -0.174867 | -0.125903 | -0.199004 |
| 6.154 | -0.152028 | -0.171782 | -0.12394 | -0.195741 |
| 6.155 | -0.149428 | -0.168671 | -0.121951 | -0.192451 |
| 6.156 | -0.146903 | -0.16561 | -0.119987 | -0.189365 |
| 6.157 | -0.144276 | -0.162474 | -0.118024 | -0.186127 |
| 6.158 | -0.141777 | -0.159363 | -0.116112 | -0.182863 |
| 6.159 | -0.139125 | -0.156277 | -0.113969 | -0.179676 |
| 6.16 | -0.136958 | -0.153523 | -0.112286 | -0.176921 |
| 6.161 | -0.134842 | -0.150897 | -0.110782 | -0.174167 |
| 6.162 | -0.132903 | -0.148423 | -0.109175 | -0.171388 |
| 6.163 | -0.130685 | -0.145823 | -0.107365 | -0.168583 |
| 6.164 | -0.128466 | -0.143298 | -0.105631 | -0.165727 |
| 6.165 | -0.12635 | -0.140799 | -0.103744 | -0.162922 |
| 6.166 | -0.124131 | -0.138274 | -0.102087 | -0.159989 |
| 6.167 | -0.121964 | -0.13575 | -0.100353 | -0.157337 |
| 6.168 | -0.119822 | -0.1332 | -0.0985675 | -0.154634 |
| 6.169 | -0.117629 | -0.130676 | -0.096757 | -0.151855 |
| 6.17 | -0.115436 | -0.128151 | -0.095074 | -0.148974 |
| 6.171 | -0.113268 | -0.125601 | -0.093289 | -0.14599 |
| 6.172 | -0.110999 | -0.123 | -0.0914785 | -0.143236 |
| 6.173 | -0.10878 | -0.120476 | -0.08977 | -0.140278 |
| 6.174 | -0.10669 | -0.117951 | -0.0880105 | -0.137549 |
| 6.175 | -0.104777 | -0.115733 | -0.0864805 | -0.135102 |
| 6.176 | -0.103196 | -0.113743 | -0.08518 | -0.13324 |
| 6.177 | -0.101513 | -0.111704 | -0.083752 | -0.13097 |
| 6.178 | -0.0999065 | -0.10974 | -0.082477 | -0.128522 |
| 6.179 | -0.0980705 | -0.107674 | -0.0810235 | -0.126304 |
| 6.18 | -0.096311 | -0.105635 | -0.0794935 | -0.123958 |
| 6.181 | -0.094475 | -0.10362 | -0.0780655 | -0.121893 |
| 6.182 | -0.092741 | -0.101606 | -0.0767395 | -0.119138 |
| 6.183 | -0.0908795 | -0.09954 | -0.075184 | -0.117303 |
| 6.184 | -0.0890945 | -0.0975 | -0.0735775 | -0.11465 |
| 6.185 | -0.087284 | -0.09546 | -0.0722005 | -0.112355 |
| 6.186 | -0.085499 | -0.093471 | -0.070747 | -0.110086 |
| 6.187 | -0.083714 | -0.0914055 | -0.0691915 | -0.107715 |
| 6.188 | -0.0819035 | -0.089391 | -0.067789 | -0.10542 |
| 6.189 | -0.0801185 | -0.087351 | -0.0663865 | -0.103176 |
| 6.19 | -0.078665 | -0.0856935 | -0.065188 | -0.101365 |
| 6.191 | -0.0773135 | -0.084087 | -0.064015 | -0.0995035 |
| 6.192 | -0.0758855 | -0.0824805 | -0.0629185 | -0.097693 |
| 6.193 | -0.0745595 | -0.080925 | -0.061873 | -0.095959 |
| 6.194 | -0.073208 | -0.079293 | -0.0607255 | -0.094123 |
| 6.195 | -0.071831 | -0.0777375 | -0.0596035 | -0.092287 |
| 6.196 | -0.0704285 | -0.076131 | -0.0583795 | -0.090349 |
| 6.197 | -0.0689495 | -0.07455 | -0.057181 | -0.0887425 |
| 6.198 | -0.067547 | -0.072969 | -0.0560335 | -0.0868555 |
| 6.199 | -0.0660935 | -0.071388 | -0.0548095 | -0.084943 |
| 6.2 | -0.0645635 | -0.069756 | -0.0535855 | -0.0830305 |
| 6.201 | -0.0631355 | -0.0681495 | -0.052387 | -0.081169 |
| 6.202 | -0.061682 | -0.0665685 | -0.0511885 | -0.079231 |
| 6.203 | -0.0602795 | -0.0649875 | -0.0500155 | -0.077344 |
| 6.204 | -0.0588515 | -0.063381 | -0.0489445 | -0.075508 |
| 6.205 | -0.0577295 | -0.062157 | -0.04795 | -0.0741055 |
| 6.206 | -0.0566075 | -0.0609075 | -0.0471085 | -0.072754 |
| 6.207 | -0.0556385 | -0.05976 | -0.046114 | -0.071275 |
| 6.208 | -0.0545675 | -0.0585105 | -0.045298 | -0.069694 |
| 6.209 | -0.05342 | -0.057261 | -0.0444565 | -0.0686485 |
| 6.21 | -0.0524 | -0.0560625 | -0.043462 | -0.067195 |
| 6.211 | -0.051278 | -0.0548385 | -0.042748 | -0.065818 |
| 6.212 | -0.050156 | -0.0536145 | -0.041728 | -0.06439 |
| 6.213 | -0.0490595 | -0.0523905 | -0.040861 | -0.0629365 |
| 6.214 | -0.0478865 | -0.051141 | -0.0398665 | -0.0615085 |
| 6.215 | -0.0467645 | -0.049917 | -0.038923 | -0.060055 |
| 6.216 | -0.045668 | -0.0487695 | -0.0379795 | -0.0585505 |
| 6.217 | -0.044495 | -0.0475455 | -0.0371125 | -0.0572245 |
| 6.218 | -0.043424 | -0.046296 | -0.0361435 | -0.0557455 |
| 6.219 | -0.042302 | -0.0451485 | -0.0352 | -0.054292 |
| 6.22 | -0.041486 | -0.0441795 | -0.034588 | -0.0531955 |
| 6.221 | -0.0406445 | -0.0432615 | -0.033874 | -0.051997 |
| 6.222 | -0.0398795 | -0.0423945 | -0.03316 | -0.0510535 |
| 6.223 | -0.0390125 | -0.041451 | -0.0325225 | -0.05011 |
| 6.224 | -0.038222 | -0.0405585 | -0.031834 | -0.0489115 |
| 6.225 | -0.037355 | -0.039615 | -0.0311455 | -0.047917 |
| 6.226 | -0.036539 | -0.0387225 | -0.030406 | -0.046795 |
| 6.227 | -0.035672 | -0.0377535 | -0.029743 | -0.0458005 |
| 6.228 | -0.034907 | -0.036912 | -0.02908 | -0.044704 |
| 6.229 | -0.033989 | -0.035943 | -0.028264 | -0.0436585 |
| 6.23 | -0.033173 | -0.035025 | -0.0275755 | -0.0424345 |
| 6.231 | -0.0323315 | -0.034158 | -0.0269635 | -0.0415165 |
| 6.232 | -0.0314135 | -0.033189 | -0.026173 | -0.0402925 |
| 6.233 | -0.0305975 | -0.032322 | -0.02551 | -0.0390685 |
| 6.234 | -0.0297305 | -0.0314295 | -0.024796 | -0.0380995 |
| 6.235 | -0.0291695 | -0.0307665 | -0.0242095 | -0.03736 |
| 6.236 | -0.028532 | -0.0300525 | -0.023776 | -0.0366715 |
| 6.237 | -0.0279455 | -0.029415 | -0.0232915 | -0.0357025 |
| 6.238 | -0.0272825 | -0.0287265 | -0.0227815 | -0.034963 |
| 6.239 | -0.0266705 | -0.0280635 | -0.022348 | -0.0340705 |
| 6.24 | -0.026033 | -0.027375 | -0.021685 | -0.0333055 |
| 6.241 | -0.0254465 | -0.0267375 | -0.021175 | -0.0325405 |
| 6.242 | -0.024809 | -0.0260235 | -0.0207415 | -0.031852 |
| 6.243 | -0.024146 | -0.025386 | -0.0202315 | -0.0309595 |
| 6.244 | -0.0236105 | -0.024723 | -0.019696 | -0.03022 |
| 6.245 | -0.022973 | -0.0240345 | -0.019186 | -0.0295315 |
| 6.246 | -0.0222845 | -0.023397 | -0.0185995 | -0.028435 |
| 6.247 | -0.021698 | -0.022683 | -0.018166 | -0.0277975 |
| 6.248 | -0.021035 | -0.0220455 | -0.017554 | -0.0265225 |
| 6.249 | -0.0204995 | -0.021459 | -0.017146 | -0.0263185 |
| 6.25 | -0.020015 | -0.020898 | -0.016738 | -0.0256045 |
| 6.251 | -0.0195305 | -0.020439 | -0.01633 | -0.0250945 |
| 6.252 | -0.0190715 | -0.01998 | -0.015973 | -0.0243295 |
| 6.253 | -0.0186635 | -0.0194955 | -0.0155395 | -0.023845 |
| 6.254 | -0.0182045 | -0.0190365 | -0.0153355 | -0.023488 |
| 6.255 | -0.0177455 | -0.0185265 | -0.0148765 | -0.0226465 |
| 6.256 | -0.0173375 | -0.018093 | -0.0145705 | -0.0222385 |
| 6.257 | -0.016853 | -0.0175575 | -0.014137 | -0.0214735 |
| 6.258 | -0.0164195 | -0.017124 | -0.0136525 | -0.0209125 |
| 6.259 | -0.0158585 | -0.0165375 | -0.013423 | -0.0205045 |
| 6.26 | -0.0155525 | -0.016155 | -0.0129385 | -0.019714 |
| 6.261 | -0.0150935 | -0.015696 | -0.0125815 | -0.0192295 |
| 6.262 | -0.0145835 | -0.015186 | -0.012199 | -0.0187705 |
| 6.263 | -0.0141245 | -0.014727 | -0.0118675 | -0.018082 |
| 6.264 | -0.0137165 | -0.014268 | -0.0115105 | -0.017521 |
| 6.265 | -0.0134105 | -0.013962 | -0.01123 | -0.017011 |
| 6.266 | -0.01313 | -0.013656 | -0.010873 | -0.0167815 |
| 6.267 | -0.012773 | -0.013248 | -0.0106945 | -0.016297 |
| 6.268 | -0.012467 | -0.012942 | -0.010414 | -0.0158125 |
| 6.269 | -0.0121355 | -0.0125595 | -0.01021 | -0.0156595 |
| 6.27 | -0.011855 | -0.0122535 | -0.009904 | -0.015073 |
| 6.271 | -0.011549 | -0.0119475 | -0.0097 | -0.014869 |
| 6.272 | -0.0112175 | -0.0115905 | -0.009343 | -0.014359 |
| 6.273 | -0.0108605 | -0.011259 | -0.0091135 | -0.0139 |
| 6.274 | -0.01058 | -0.0109275 | -0.008935 | -0.0136705 |
| 6.275 | -0.010274 | -0.010596 | -0.008527 | -0.013135 |
| 6.276 | -0.0099425 | -0.0102645 | -0.008221 | -0.0125995 |
| 6.277 | -0.0095345 | -0.009882 | -0.008068 | -0.012421 |
| 6.278 | -0.009305 | -0.009627 | -0.007762 | -0.0118855 |
| 6.279 | -0.008999 | -0.009321 | -0.007609 | -0.011707 |
| 6.28 | -0.008795 | -0.0090915 | -0.0072775 | -0.0111715 |
| 6.281 | -0.008591 | -0.008862 | -0.007252 | -0.0109165 |
| 6.282 | -0.008387 | -0.0086325 | -0.006997 | -0.0106615 |
| 6.283 | -0.0081065 | -0.0083775 | -0.006793 | -0.0104065 |
| 6.284 | -0.0079025 | -0.008148 | -0.00664 | -0.0102025 |
| 6.285 | -0.0076985 | -0.007944 | -0.006487 | -0.0099475 |
| 6.286 | -0.007418 | -0.0076635 | -0.00613 | -0.0095905 |
| 6.287 | -0.007265 | -0.007485 | -0.0061045 | -0.0092335 |
| 6.288 | -0.007061 | -0.007281 | -0.005926 | -0.0086725 |
| 6.289 | -0.0067805 | -0.0070005 | -0.005773 | -0.0087745 |
| 6.29 | -0.0065255 | -0.0067455 | -0.0056455 | -0.0085195 |
| 6.291 | -0.0063725 | -0.006567 | -0.0052885 | -0.0081625 |
| 6.292 | -0.006092 | -0.0062865 | -0.005161 | -0.0079585 |
| 6.293 | -0.0059135 | -0.0060825 | -0.0049825 | -0.0077035 |
| 6.294 | -0.0057605 | -0.005904 | -0.004906 | -0.007474 |
| 6.295 | -0.005582 | -0.0057255 | -0.0047275 | -0.0072445 |
| 6.296 | -0.00548 | -0.005649 | -0.0045745 | -0.0069385 |
| 6.297 | -0.005327 | -0.005496 | -0.004498 | -0.006709 |
| 6.298 | -0.0051995 | -0.005343 | -0.004345 | -0.0065305 |
| 6.299 | -0.0049955 | -0.005139 | -0.004141 | -0.006454 |

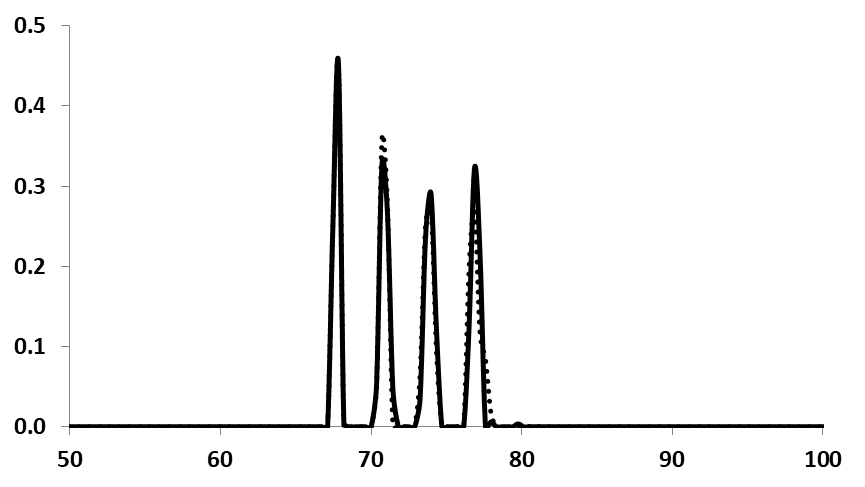

**6.4-7.0 cm**

**6.4-6.9 cm**

**Sedimentation coefficient, S**

**c(s)**

**B)**

|  |  |  |
| --- | --- | --- |
| 40 | 0 | 0.000201884 |
| 40.3614 | 0 | 0 |
| 40.7229 | 0 | 0 |
| 41.0843 | 0 | 0 |
| 41.4458 | 0 | 0 |
| 41.8072 | 0 | 0 |
| 42.1687 | 0 | 0 |
| 42.5301 | 0 | 0 |
| 42.8916 | 0 | 0 |
| 43.253 | 0.000355237 | 0.000136039 |
| 43.6145 | 7.27689e-05 | 0.000270857 |
| 43.9759 | 0 | 0 |
| 44.3373 | 0 | 0 |
| 44.6988 | 0 | 0 |
| 45.0602 | 0 | 0 |
| 45.4217 | 0 | 0 |
| 45.7831 | 0 | 0 |
| 46.1446 | 0 | 0 |
| 46.506 | 0 | 0 |
| 46.8675 | 0 | 0 |
| 47.2289 | 0 | 0 |
| 47.5904 | 0 | 0 |
| 47.9518 | 0 | 0 |
| 48.3133 | 0 | 0 |
| 48.6747 | 0 | 0 |
| 49.0361 | 0 | 0 |
| 49.3976 | 0 | 0 |
| 49.759 | 0 | 0 |
| 50.1205 | 0 | 0 |
| 50.4819 | 0 | 0 |
| 50.8434 | 0 | 0 |
| 51.2048 | 0 | 0 |
| 51.5663 | 0 | 0 |
| 51.9277 | 0 | 0 |
| 52.2892 | 0 | 0 |
| 52.6506 | 0 | 0 |
| 53.012 | 0 | 0 |
| 53.3735 | 0 | 0 |
| 53.7349 | 0 | 0 |
| 54.0964 | 0 | 0 |
| 54.4578 | 0 | 0 |
| 54.8193 | 0 | 0 |
| 55.1807 | 0 | 0 |
| 55.5422 | 0 | 0 |
| 55.9036 | 0 | 0 |
| 56.2651 | 0 | 0 |
| 56.6265 | 0 | 0 |
| 56.988 | 0 | 0 |
| 57.3494 | 0 | 0 |
| 57.7108 | 0 | 0 |
| 58.0723 | 0 | 0 |
| 58.4337 | 0 | 0 |
| 58.7952 | 0 | 0 |
| 59.1566 | 0 | 0 |
| 59.5181 | 0 | 0 |
| 59.8795 | 0 | 0 |
| 60.241 | 0 | 0 |
| 60.6024 | 0 | 0 |
| 60.9639 | 0 | 0 |
| 61.3253 | 0 | 0 |
| 61.6867 | 0 | 0 |
| 62.0482 | 0 | 0 |
| 62.4096 | 0 | 0 |
| 62.7711 | 0 | 0 |
| 63.1325 | 0 | 0 |
| 63.494 | 0 | 0 |
| 63.8554 | 0 | 0 |
| 64.2169 | 0 | 0 |
| 64.5783 | 0 | 0 |
| 64.9398 | 0 | 0 |
| 65.3012 | 0 | 0 |
| 65.6627 | 0 | 0 |
| 66.0241 | 0 | 0 |
| 66.3855 | 0 | 0 |
| 66.747 | 0 | 0 |
| 67.1084 | 0 | 0 |
| 67.4699 | 0.258218 | 0.255938 |
| 67.8313 | 0.44958 | 0.453087 |
| 68.1928 | 0 | 0.00462097 |
| 68.5542 | 0 | 0 |
| 68.9157 | 0 | 0 |
| 69.2771 | 0 | 0 |
| 69.6386 | 0 | 0 |
| 70 | 0 | 0 |
| 70.3614 | 0.048484 | 0.0532153 |
| 70.7229 | 0.359484 | 0.328024 |
| 71.0843 | 0.27725 | 0.265456 |
| 71.4458 | 0 | 0.0472645 |
| 71.8072 | 0 | 0 |
| 72.1687 | 0 | 0 |
| 72.5301 | 0 | 0 |
| 72.8916 | 0 | 0 |
| 73.253 | 0.0662713 | 0.036133 |
| 73.6145 | 0.252414 | 0.23682 |
| 73.9759 | 0.273455 | 0.290696 |
| 74.3373 | 0.0947033 | 0.1215 |
| 74.6988 | 0 | 0 |
| 75.0602 | 0 | 0 |
| 75.4217 | 0 | 0 |
| 75.7831 | 0 | 0 |
| 76.1446 | 0 | 0 |
| 76.506 | 0.199035 | 0.126046 |
| 76.8675 | 0.27089 | 0.323112 |
| 77.2289 | 0.11665 | 0.215882 |
| 77.5904 | 0.0836988 | 0 |
| 77.9518 | 0.0168964 | 0.00620387 |
| 78.3133 | 0 | 0 |
| 78.6747 | 0 | 0 |
| 79.0361 | 0 | 0 |
| 79.3976 | 0 | 0 |
| 79.759 | 0 | 0.00361225 |
| 80.1205 | 0 | 0 |
| 80.4819 | 0.0004031 | 0 |
| 80.8434 | 0 | 0 |
| 81.2048 | 0.00045999 | 0 |
| 81.5663 | 0 | 0 |
| 81.9277 | 0 | 0 |
| 82.2892 | 0 | 0 |
| 82.6506 | 0 | 0 |
| 83.012 | 0 | 0 |
| 83.3735 | 0 | 0 |
| 83.7349 | 0 | 0 |
| 84.0964 | 0 | 0 |
| 84.4578 | 0 | 0 |
| 84.8193 | 0 | 0 |
| 85.1807 | 0 | 0 |
| 85.5422 | 0 | 0 |
| 85.9036 | 0 | 0 |
| 86.2651 | 0 | 0 |
| 86.6265 | 0 | 0 |
| 86.988 | 0 | 0 |
| 87.3494 | 0 | 0 |
| 87.7108 | 0 | 0 |
| 88.0723 | 0 | 0 |
| 88.4337 | 0 | 0 |
| 88.7952 | 0 | 0 |
| 89.1566 | 0 | 0 |
| 89.5181 | 0 | 0 |
| 89.8795 | 0 | 0 |
| 90.241 | 0 | 0 |
| 90.6024 | 0 | 0 |
| 90.9639 | 0 | 0 |
| 91.3253 | 0 | 0 |
| 91.6867 | 0 | 0 |
| 92.0482 | 0 | 0 |
| 92.4096 | 0 | 0 |
| 92.7711 | 0 | 0 |
| 93.1325 | 0 | 0 |
| 93.494 | 0 | 0 |
| 93.8554 | 0 | 0 |
| 94.2169 | 0 | 0 |
| 94.5783 | 0 | 0 |
| 94.9398 | 0 | 0 |
| 95.3012 | 0 | 0 |
| 95.6627 | 0 | 0 |
| 96.0241 | 0 | 0 |
| 96.3855 | 0 | 0 |
| 96.747 | 0 | 0 |
| 97.1084 | 0 | 0 |
| 97.4699 | 0 | 0 |
| 97.8313 | 0 | 0 |
| 98.1928 | 0 | 0 |
| 98.5542 | 0 | 0 |
| 98.9157 | 0 | 0 |
| 99.2771 | 0 | 0 |
| 99.6386 | 0 | 0 |
| 100 | 0 | 0 |
| 100.361 | 0 | 0 |
| 100.723 | 0 | 0 |
| 101.084 | 0 | 0 |
| 101.446 | 0 | 0 |
| 101.807 | 6.42906e-05 | 0 |
| 102.169 | 0 | 0 |
| 102.53 | 0 | 0 |
| 102.892 | 0 | 0 |
| 103.253 | 0 | 0 |
| 103.614 | 0 | 0 |
| 103.976 | 0.000207083 | 0.000144543 |
| 104.337 | 0 | 0 |
| 104.699 | 0 | 0 |
| 105.06 | 0 | 0 |
| 105.422 | 0 | 0 |
| 105.783 | 0 | 0 |
| 106.145 | 0 | 0 |
| 106.506 | 0 | 0 |
| 106.867 | 0 | 0 |
| 107.229 | 0 | 0 |
| 107.59 | 0 | 0 |
| 107.952 | 0 | 0 |
| 108.313 | 0 | 0 |
| 108.675 | 0 | 0 |
| 109.036 | 0 | 0 |
| 109.398 | 0 | 0 |
| 109.759 | 0 | 0 |
| 110.12 | 0 | 0 |
| 110.482 | 0 | 0 |
| 110.843 | 0 | 0 |
| 111.205 | 0 | 0 |
| 111.566 | 0 | 0 |
| 111.928 | 0 | 0 |

**Figure 6S**

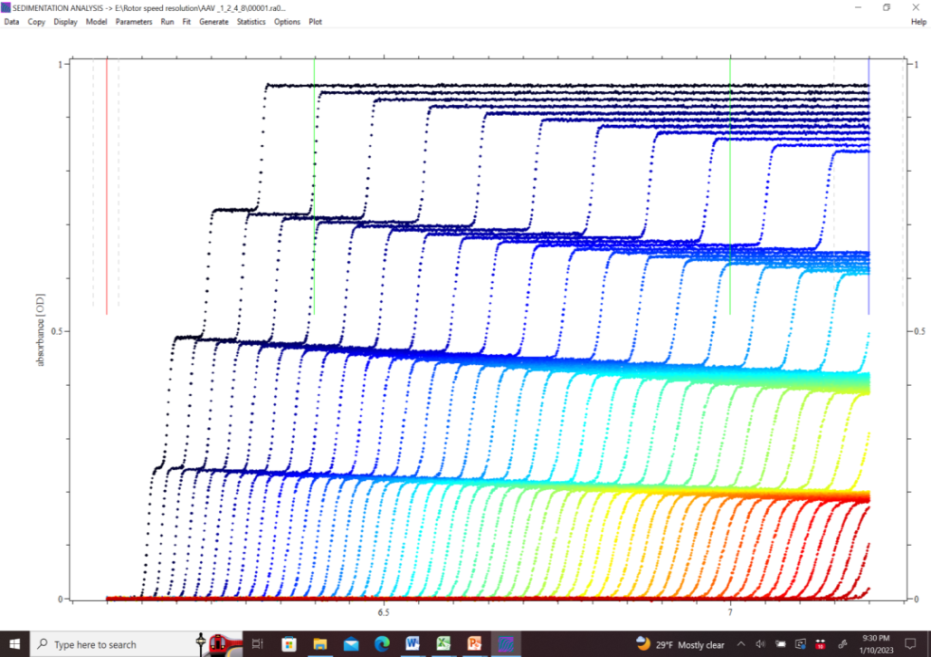

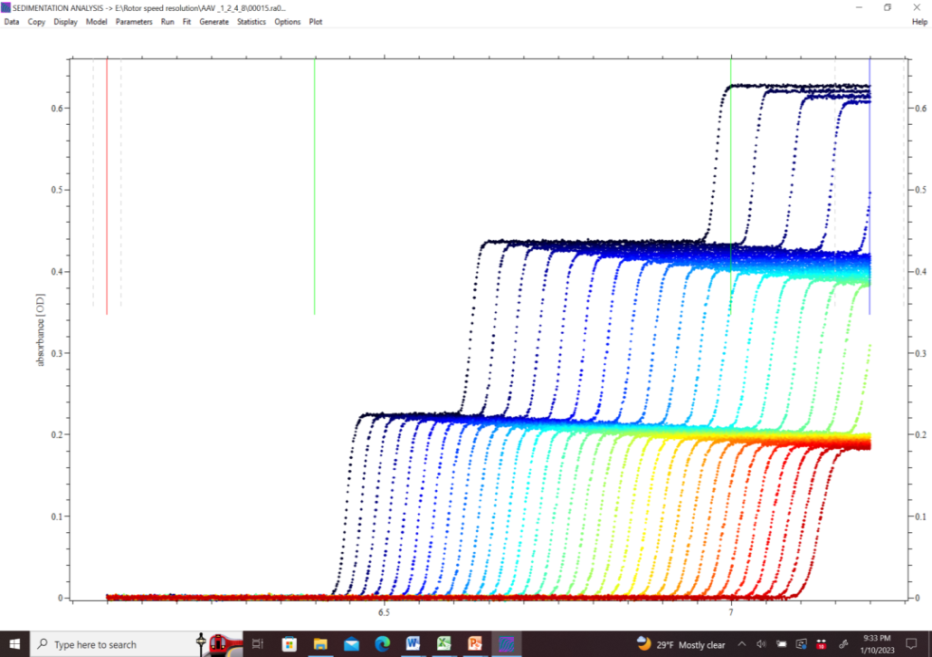

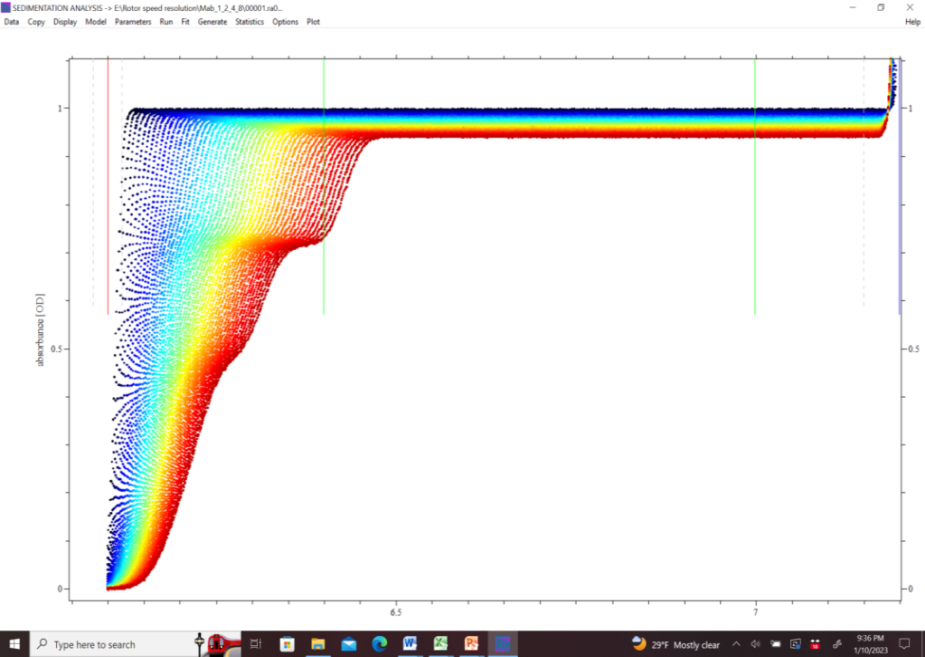

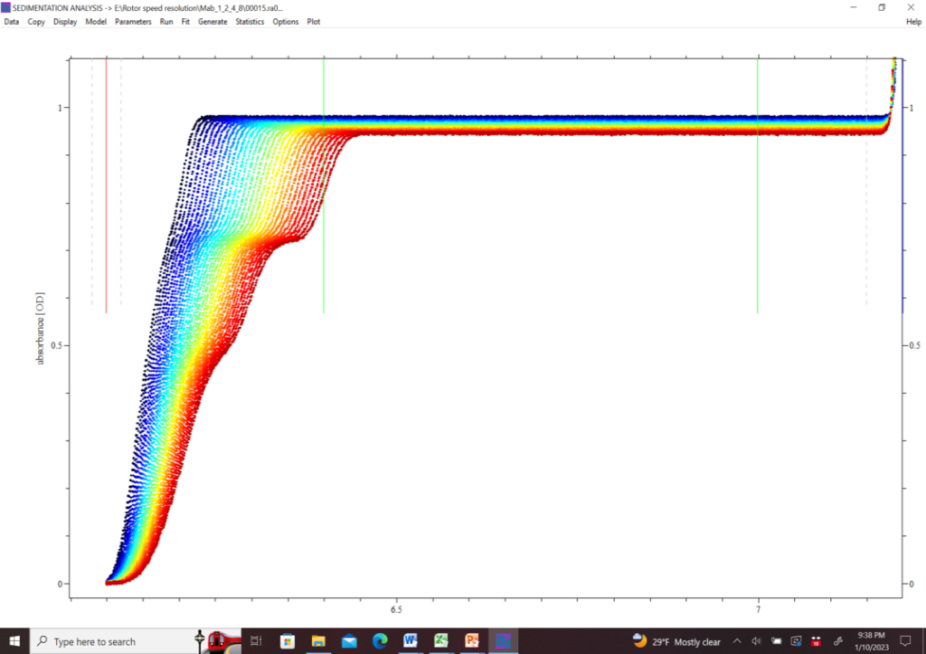

**A)**

**B)**

**C)**

**1**

**2**

**3**

**4**

**2**

**3**

**1**

**4**

**3**

**1 & 2**

**4**

**3**

**1 & 2**

**Radius, cm**

**Concentration, arbitrary units**

**Radius, cm**

**Concentration, arbitrary units**

**Concentration, arbitrary units**

**Concentration, arbitrary units**

**D)**

**Radius, cm**

**Radius, cm**

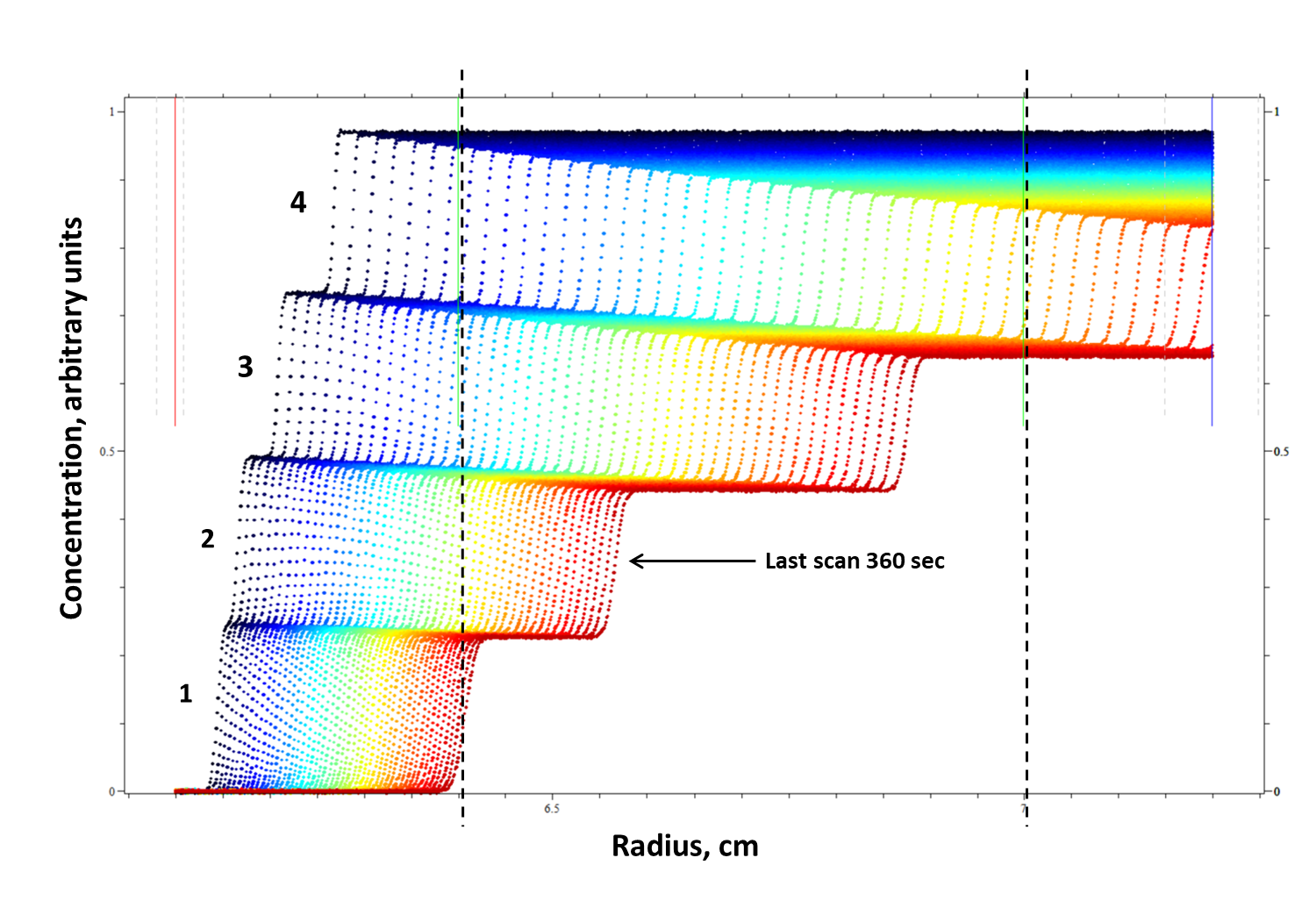

**Figure 7S**

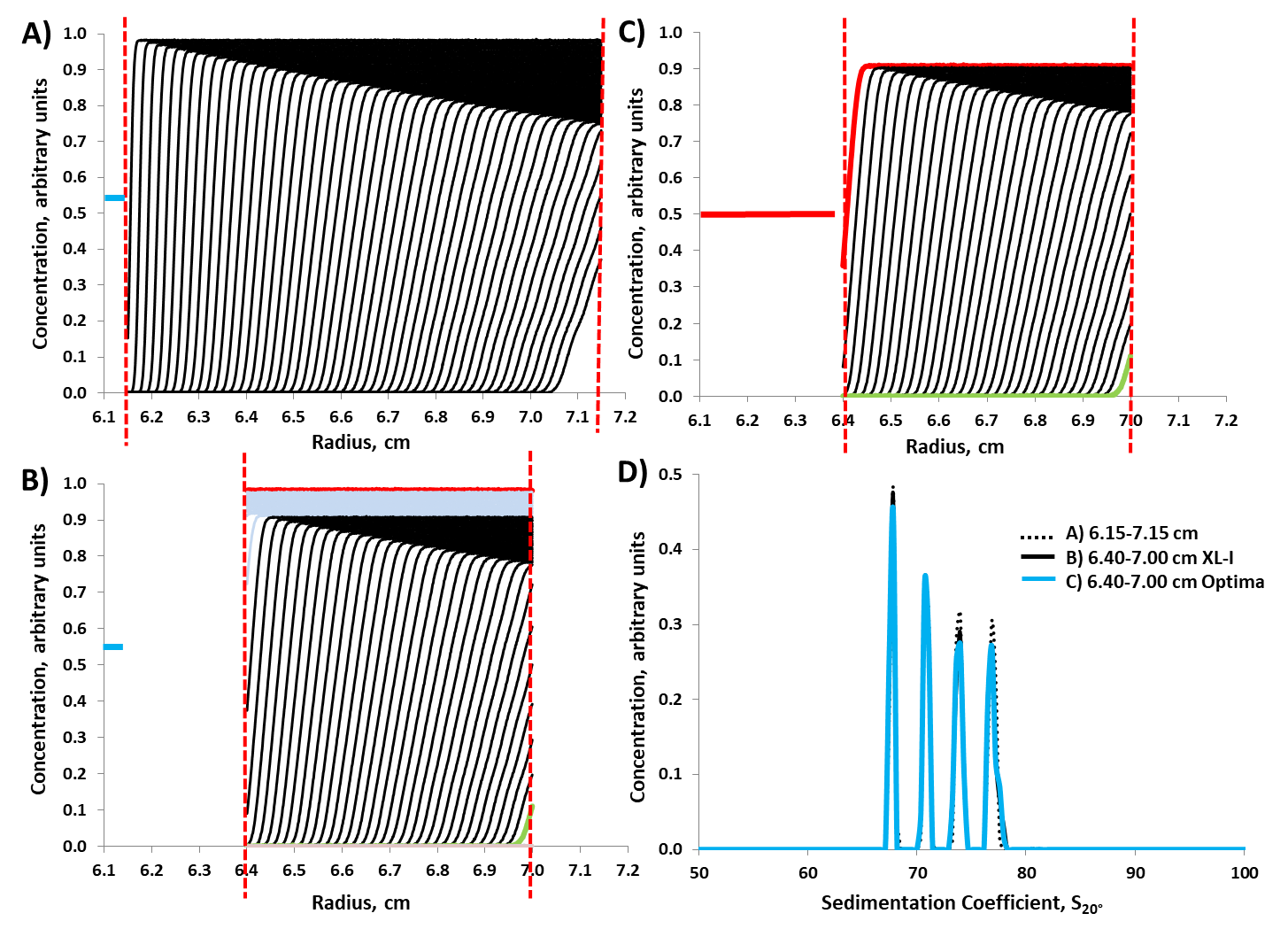

**Figure 8S**

**Supplemental Material Figure Legends**

**Figure 1S.** An outline of the process steps, given in the Material and Methods section 2.3.2, for generating a “*solvent mismatch interference difference signal plot*” from separate simulated reference and sample sector solvent SV-AUC data generated using the following specified (upper limit) solvent mismatch parameters of 0.1 cm difference in menisci position and 5% difference in composition between the reference and sample sectors (as described in Material and Methods section 2.3.2), which in this case is specific for 0.15 M NaCl at 45K rpm at 10 °C using a data scan time interval of 19 sec.

**Figure 2S.** A pictorial version of the process steps given in the Material and Methods section 2.3.2 and outlined in Figure 1S, for generating the “*solvent mismatch interference difference signal plot*” from simulated 0.15 M NaCl (solvent) hs-SV-AUC at 45K rpm at 10 °C data using a data scan interval of 19 sec generated with specifically defined solvent mismatches between the reference and sample sectors (see Material and Methods section 2.3.2): **A)** Resulting 1^st^ and last reference and sample sector solvent SV-AUC scans (at an arbitrary concentration of 1.0 unit for the reference sector over the radial range 6.0-7.2 cm and at an arbitrary concentration of 0.95 unit for the sample sector over the radial range 6.1-7.2 cm) corresponding to the time points for the 1^st^ and last simulated AAV hs-SV-AUC data scan used in AAV c(s) data analysis (see Figure 4A, which shows the last AAV scan corresponds to scan 46), **B)** 1^st^ and last reference and sample sector solvent SV-AUC scans over the radial range of 6.1-7.2 cm where arbitrary concentration units were converted to units of fringes by multiplying all scan radial readings by 25.5 fringes, **C)** the resulting 1^st^ and last solvent interference contribution scans, in fringes, computed by subtracting the 1^st^ solvent reference sector scan from the 1^st^ solvent sample sector scan and the last solvent reference sector scan from the last solvent sample sector scan, and **D)** the resulting “*solvent mismatch interference difference signal plo*t” computed by subtracting the 1^st^ solvent interference contributing scan from the last solvent interference contributing scan.

**Figure 3S. A)** Overlay of actual experimental AAV-hs-SV-AUC processed interference data scans obtained on an Optima at 45K rpm at 10 °C using a data scan time interval of 6 sec. The occasional erratic radial spacing of the overlay data scans in this plot appears to be a characteristic feature of the rapid data acquisition rate (6 sec) on the Optima. Nevertheless the reported data scan time stamp associated with each scan is valid. **B)** The overlay of residual plots for each data scan shown in “**A**” resulting from AAV c(s) data analysis. **C)** Distribution plot of the resulting fit residuals shown in “**B**” relative to their anticipated normal distribution based on the data’s RMSD (outlined in red), which in this example had a value of 0.0012 fringe, yielding a residual histogram H value of 0.3%.

**Figure 4S. A)** The overlay of the four area normalized c(s) vs S distribution plots shown in Figure 6, **B)** same plot as in “**A**”, but expanded in the X-axis.

**Figure 5S.** An important feature in the successful operation of AAV hs-SV-AUC (45K rpm and 10 °C) is that the volume of AAV sample solution placed in the sample sector be sufficient so that it’s meniscus is located at a radial position of about 6.1 cm or less, and that the references sector solution is filled so its meniscus is at a radial position value equal or less than the sample sector’s meniscus position, but the difference in menisci positions should not be more than 0.1 cm. In addition, the solvent composition of the sample and reference solution must also not differ by more than ± 5%. Nevertheless, “*normalized* *solvent mismatch interference difference signal plot”* (see Material and Methods section 2.4) shown in **A)** indicates that if these two upper limits of mismatch are greatly exceeded the resulting impact will amount to a reduction in the radial-time window’s flatness of at most about a 0.1 cm (see black dashed outline box). **B**) Shows that this small radial reduction (0.1 cm) in the radial-time window has a minimum impact on the hs-SV-AUC method, as shown by the high similarity in the overlay plot of the c(s) vs S distribution obtained for the simulated AAV sample shown in Table 1 using a radial-time window of 6.4-7.0 cm vs 6.4-6.9 cm, where the time window in both cases was the same, ~1000 sec. Overall, the results provided in this figure supports hs-SV-AUC’s robustness in tolerating significant departures from the paper’s stated upper solvent mismatch limits.

**Figure 6S.** Simulated hs-SV-AUC (45K rpm and 10 °C) data generated under the same experimental protocol conditions indicated in Figure 4A for the full radial-time window (6.15-7.15 cm) to assess the upper S value limit that can be monitored using a 2^nd^ hypothetical AAV sample [consisting of equal amounts of the **1** - full monomeric AAV (5.23 MDa, 76.6 S) plus three of its unique aggregates: **2** - dimer (10.5 MDa, 114.9 S), **3** - tetramer (20.9 MDa, 186.9 S in a tetrahedron configuration) and **4** - octamer (41.8 MDa, 281.9 S in a cube configuration)] for **A**) the XL-I’s interferometer and **B**) the Optima’s interferometer. A similar type of analysis to assess the lower S value limit that can be monitored using a hypothetical monoclonal antibody, mAb, sample [consisting of a **1** - monomeric mAb (150 KDa, 5.0 S) plus three of its unique aggregates: **2** - dimer (300 kDa, 7.5 S), **3** - tetramer (600 kDa, 12.2 S in tetrahedron configuration) and **4** - hexamer (1,200 kDa, 16.2 S in octahedron configuration)] for **C**) the XL-I’s interferometer and **D**) the Optima’s interferometer. Note S values assigned to the AAV aggregates of the full AAV monomeric particle and mAb aggregates of the monomeric mAb were calculated using known S value ratios for specifically configured aggregates of a spherical monomer particle [1]. All above cited S values are S_10°_ values. Vertical black dashed lines define the radial window (6.4-7.0 cm) of data that would be used in c(s) analysis of AAV hs-SV-AUC (at 45K rpm at 10 °C) data, where solvent contribution to the AAV signal can easily be removed.

**Figure 7S.** Opportunity to expand the upper S_10_ value of the AAV hs-SV-AUC protocol for the XL-I by reducing data scan time interval from 19 sec to 5 sec. Using the same hypothetical AAV sample and experimental conditions used in Figure 6SA the resulting reduction in data scan time interval shows that particles 2, 3 and 4 can now be adequately monitored (relative to the situation seen in Figure 6SA), significantly increasing the upper size limit that can be detected when doing AAV hs-SV-AUC at 45K rpm at 10 °C from about 150 to 300 S_10°_. However, now particle “1” would not be detected if only 50 scans were acquired. Nevertheless, if this faster data acquisition was maintained for the normal time period used for collecting 50 scans at the data acquisition rate of 19 sec (about 1000 sec), particle 1 (along with particles down to about 20 S_10°_) could also be monitored. To achieve this goal all one needs to do is to increases the number of data scans that are collected (using the data scan time interval of 5 sec) to about 200 scans. However, this opportunity can only be exploited on using the XL-I. It is not available to the Optima due to the required time delay in setting up its interferometer’s laser. Vertical black dashed lines shown in the plot define the radial window (6.4-7.0 cm) used in c(s) analysis on AAV hs-SV-AUC data, where solvent interference signal contributions can easily be removed.

**Figure 8S.** A reassessment of the comparison of full vs restricted radial-time window data shown in Figure 4 taking into account the automated operating procedure used in setting up the Optima’s interference’s laser timing relative to the manual operating procedure used for the XL-I’s interferometer. **A)** Corresponds to Figure 4A showing the 50 AAV hs-SV-AUC data scans that the XL-I’s interferometer would record given the stated experimental conditions, blue horizontal line corresponds to a small loss in data scans resulting from the time (amounting to about 120 sec) needed for the rotor to reach its running speed of 45K rpm at 10 °C (given an acceleration rate of 400 rpm/sec). **B)** Same as “**A**”, but in this case what is being shown is the “*regions*” of the AAV hs-SV-AUC data scans that the XL-I can use in doing AAV c(s) analysis under the restricted radial-time window of 6.4-7.0 cm and about 1000 sec (which includes red region of the 1^st^ scans, light blue region of scans 2-13, and light green region of scan 46, which is the last data scan used in AAV c(s) analysis). Red dashed vertical lines correspond to radial position, 6.4 and 7.0 cm used in AAV hs-SV-AUC c(s) analysis. **C)** Same as “**A**”, but in this case what is being shown is the “*regions*” of the AAV hs-SV-AUC data scans that the Optima can use when doing AAV hs-SV-AUC c(s) analysis due to the impact of the automated procedure used to set up its interferometer’s laser, which adds an additional time delay (which can vary a bit from run to run) to the required time delay for the rotor to reach run speed (amounting to a total delay of about 400 sec, which is indicated by the horizontal solid red line). As a result of this extended time delay, region of data scans are lost relative to the XL-I. These scan regions correspond to the red and light blue scans indicated in “**B**”. The data scan now shown in “C” includes a red scan corresponding to the 1^st^ scan acquired by the Optima, and is equivalent (approximately) in time to scan 14 in “**B**”, while the light green scan still corresponds to the Optima scan 32 which is equivalent (approximately) in time to scan 46 in “**B**”. The red dashed vertical lines correspond to the radial window, 6.4 and 7.0 cm used in AAV hs-SV-AUC c(s) analysis. **D)** Overlay of the resulting computed AAV c(s) vs S distribution plots from data contained within the vertical dash lines in “**B**” and “**C**” indicates little difference in the resulting computed AAV c(s) vs S distribution plots given the difference in data used in each c(s) analysis.

**Supplemental Material References**

1. J. Gracia De La Torre, Sedimentation coefficient of complex biological particles, in: A.J. Harding, A.J. Rowe, J.C. Horton, Eds., Royal Society of Chemistry, Redwood Press, Cambridge, UK, 1992, pp.333-345.
